# BBSome trains remove activated GPCRs from cilia by enabling passage through the transition zone

**DOI:** 10.1101/180620

**Authors:** Fan Ye, Andrew R. Nager, Maxence V. Nachury

**Affiliations:** Department of Ophthalmology, University of California San Francisco, CA 94143, USA; Department of Molecular and Cellular Physiology, Stanford University School of Medicine, CA 94305, USA

## Abstract

A diffusion barrier at the transition zone enables the compartmentalization of signaling molecules by cilia. The BBSome and the small GTPase Arl6, which triggers BBSome coat polymerization, are required for the exit of activated signaling receptors from cilia, but how diffusion barriers are crossed when membrane proteins exit cilia remains to be determined. Here we found that activation of the ciliary GPCRs Smoothened and SSTR3 drove the Arl6-dependent assembly of large, highly processive and cargo-laden retrograde BBSome trains. Single-molecule imaging revealed that the assembly of BBSome trains enables the lateral transport of ciliary GPCRs across the transition zone. Yet, the removal of activated GPCRs from cilia was inefficient because a second, periciliary diffusion barrier was infrequently crossed. We conclude that exit from cilia is a two-step process in which BBSome/Arl6 trains first moves activated GPCRs through the transition zone before a periciliary barrier can be crossed.

**Summary:** Upon activation, GPCRs must exit cilia for appropriate signal transduction. Using bulk imaging of BBSome and single molecule imaging of GPCRs, Ye et al. demonstrate that retrograde BBSome trains assemble on-demand upon GPCR activation and ferry GPCRs across the transition zone. Yet, ciliary exit often fails because of a second diffusion barrier.

## Introduction

Diffusion barriers establish the identity of the apical membrane in polarized epithelial cells, of the axon in neurons, of the daughter cell in budding yeast, and of cilia by impeding the lateral movement of membrane proteins (Trimble and Grinstein, 2015). The compartmentalization of cilia enables dynamic changes in ciliary composition through regulated trafficking. Upon Hedgehog pathway activation, the 7-transmembrane protein Smoothened accumulates in cilia while ciliary exit of the G protein coupled receptor (GPCR) GPR161 ensures the appropriate transduction of Hedgehog signals (Bangs and Anderson, 2017; Nager et al., 2017). While trafficking across the tight junction, the axon initial segment and the yeast bud neck involves a vesicular carrier intermediate, the mechanisms of ciliary barrier crossing remain undetermined. The ciliary diffusion barrier has been localized to the transition zone, an ultrastructural specialization between the transition fibers of the basal body and the cilium shaft (Garcia-Gonzalo and Reiter, 2012; Gonçalves and Pelletier, 2017). Three hypotheses have been advanced for crossing the transition zone (Nachury et al., 2010; Jensen and Leroux, 2017). First, the detection of vesicles inside the transition zone indicates that a vesicular carrier may transport cargoes across this barrier (Jensen et al., 2004; Chuang et al., 2015). Second, indirect evidence for lateral transport between plasma and ciliary membranes (Hunnicutt et al., 1990; Milenkovic et al., 2009) suggests that membranous cargoes laterally traverse the transition zone by active transport. Third, the regulated opening of a gate inside the transition zone may let selective cargoes move through this membranous barrier (Dyson et al., 2017).

The active transport of proteins inside cilia-termed intraflagellar transport-is powered by microtubule motors moving along axonemal microtubules. It is now clear that axonemal precursors such as α/β-tubulin are delivered to the tip of cilia by anterograde IFT trains (Lechtreck, 2015; Kubo et al., 2016). In contrast, it is not known where, when and how membrane proteins are selected for ciliary exit and prior studies of ciliary signaling receptors dynamics by single-molecule imaging failed to uncover extended IFT movements (Ye et al., 2013; Milenkovic et al., 2015).

## Results

### Low-level expression recapitulates physiological ciliary trafficking dynamics

To characterize transition zone crossing by membranous cargoes, we sought a system where membrane proteins move across the transition zone in a synchronized manner. GPR161 and the prototypical ciliary GPCR somatostatin receptor 3 (SSTR3) both undergo retrieval from the cilium and back into the cell upon activation (Mukhopadhyay et al., 2013; Green et al., 2016). GPR161, a core component of the Hedgehog pathway that couples to Gα_s_, exits cilia when Smoothened is activated either indirectly by Hedgehog or directly by the Smoothened agonist SAG (Pal et al., 2016) (Fig. 1A). Meanwhile, SSTR3 is a well-characterized Gαi-coupled receptor that has undergoes agonist-dependent retrieval (Nager et al., 2017; Green et al., 2016) (Fig. **1A**). Consistent with previous studies on SSTR3 in hippocampal neurons (Green et al., 2015), most of SSTR3 immunofluorescence was lost from neuronal cilia after 6 h treatment with the ligand somatostatin-14 (sst) or the SSTR3-specific agonist L796,778 (Fig. **1B** and S1A-C). Similar to endogenous SSTR3, ciliary exit of endogenous GPR161 proceeds over the course of several hours (Mukhopadhyay et al., 2013). Signal-dependent retrieval is thus a considerably slower process than signal-dependent endocytosis.

**Figure 1.**
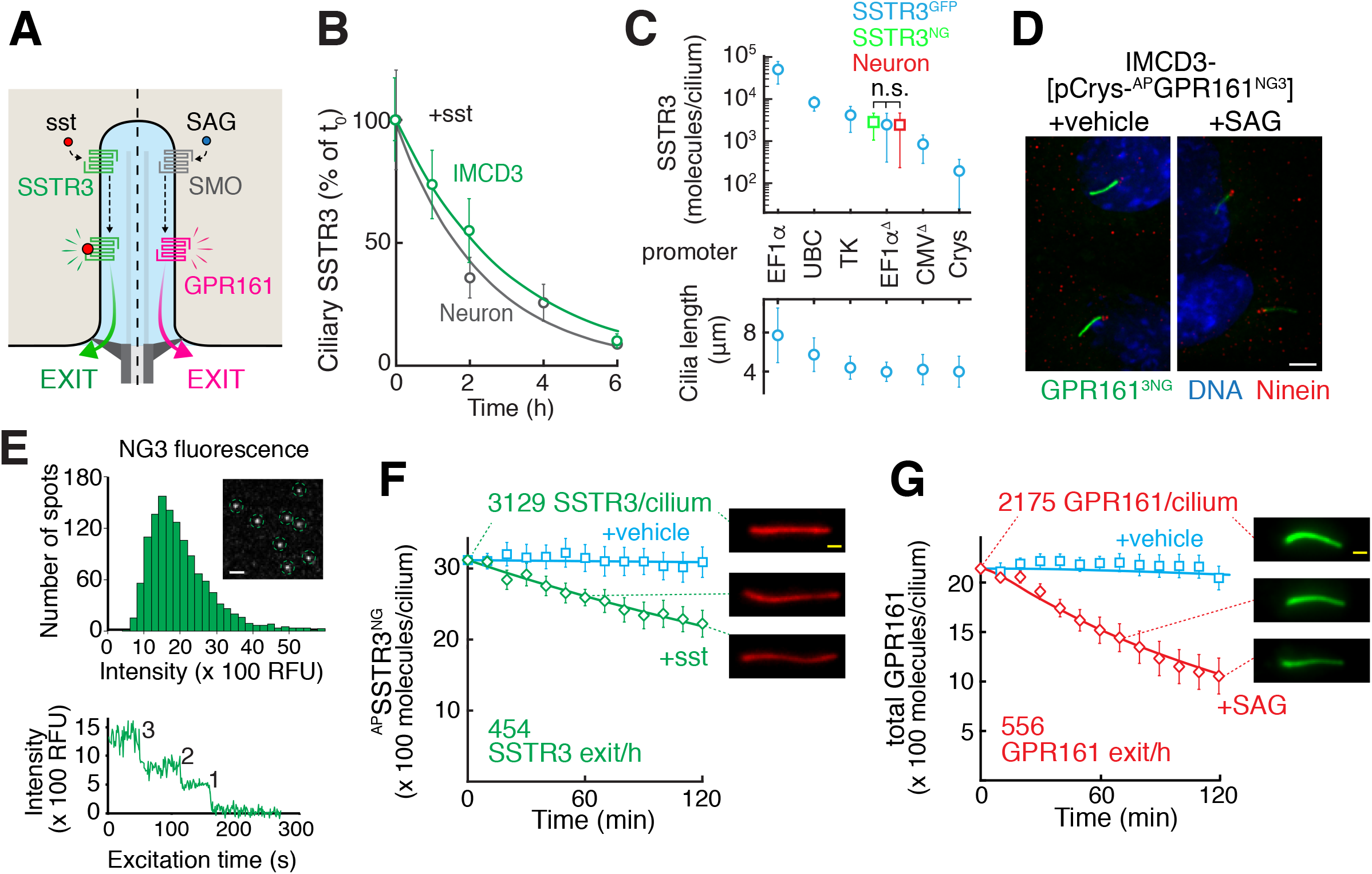
Reconstitution of signal-dependent retrieval of SSTR3 and GPR161 (**A**) Diagram of the signal-dependent retrieval systems under study. Left: addition of somatostatin triggers SSTR3 exit from cilia by directly activating SSTR3. Right: addition of Smoothened agonist (SAG) activates the Hedgehog pathway and promotes GPR161 retrieval. (**B**) Kinetics of SSTR3 disappearance from cilia of primary hippocampal neurons and of IMCD3 stably expressing ^AP^SSTR3^NG^ under the control of the TATA-less EF1α promoter were estimated by quantitation of immunofluorescence signals following addition of sst. The entire data set for the sst condition is shown in Fig. **S1B**. Data were fitted to a single exponential. Error bars: 95% CI. ***N*** = 280-424 cilia (neurons), 57-80 cilia (IMCD3). (**C**) High level expression of SSTR3 drives elongation of primary cilia. Top: ^AP^SSTR3^GFP^ driven by various promoters or ^AP^SSTR3^NG^ driven by EF1α^Δ^ promoter was expressed stably at the FlpIn locus of IMCD3 cells and ciliary fluorescence levels were measured and compared to a GFP calibrator (Breslow et al., 2013) or a NG calibrator (see Methods). Endogenous SSTR3 levels were estimated by comparative immunostaining (see Methods). A Mann-Whitney test was used for pairwise comparisons of the number of SSTR3 molecules per cilia in neurons and in IMCD3 cells expressing ^AP^SSTR3^NG^ or ^AP^SSTR3^GFP^ under the control of pEF1α^Δ^ n.s.: no significant differences were observed. ***P*** > 0.05. ***N*** = 10-38 cilia. Error bars: SD. Bottom: Effect of ^AP^SSTR3^GFP^ expression on cilium length. Cilia lengths were measured in the GFP channel by live-cell imaging. Error bars represent SD, ***N*** = 10-38 cilia. Cilium lengthening upon GPCR overexpression was previously reported by (Guadiana et al., 2013). (**D**) IMCD3-[pCrys-^AP^GPR161^3NG^] were treated for 2 h with either SAG or vehicle. ^AP^GPR161^3NG^ was visualized by NG fluorescence, basal bodies of cilia stained with Ninein. All cells were pre-treated with the translation inhibitor emetine to eliminate signals from new protein synthesis. Scale bar: 4 μm. (**E**) Absolute quantitation of ciliary GPCR abundance. Top: Calibration of single molecule fluorescence intensity. Bacterially expressed triple NeonGreen (NG3) protein was spotted on glass coverslips (inset) and the fluorescent intensity of each individual NG3 was measured. ***N*** = 1257 particles measured. Bottom: The 3-step photobleaching of a representative spot shows that the fluorescence was emitted by a single NG3 molecule. The measured fluorescence intensity of NG3 was used to calibrate NG- and NG3-tagged SSTR3, GPR161, BBS5 and IFT88. (**F**) IMCD3-[pEF1α^Δ^-^AP^SSTR3^NG^] cells were treated with vehicle or sst for 2h. Stable expression of an ER-localized biotin ligase BirA enables the biotinlylation of ^AP^SSTR3 with the biotin exists in the DMEM/F12 cell culture medium. Ciliary ^AP^SSTR3 was pulse-labeled by Alexa647-conjugated mSA (mSA647) for 5-10 min before imaging (see Methods for details). The absolute number of ^AP^SSTR3^NG^ molecules per cilia at t¿ was calculated by measuring the NG signal and using the NG3 calibrator. For all other time points, the ratio in ciliary mSA647 signal compared to t_0_ was used to calculate the absolute number of molecules (see Methods for details). Data were fitted to a single exponential. Error bars: 95% CI. ***N*** = 14 cilia. (**G**) IMCD3-[pCrys-GPR161^3NG^] cells were treated with SAG or vehicle for 2h. NG fluorescence was tracked in individual cilia and the ratio of GPR161^3NG^ to endogenous GPR161 was used to calculate the total levels of GPR161 as detailed in Method. Data were fitted to a single exponential. Error bars: 95% CI. ***N***=12-20 cilia.

To dissect ciliary exit, we expressed GPCRs in mouse Inner Medullar Collecting Duct (IMCD3) kidney cells, a widely used cell line for ciliary trafficking studies. GPCRs were tagged on the intracellular C-terminus with a fluorescent protein (GFP or NeonGreen, NG, (Shaner et al., 2013)) while a biotinylation acceptor peptide (AP) on the extracellular N-terminus combined with co-expression of the biotin ligase BirA enabled pulse-chase studies with fluorescently labeled monovalent streptavidin (mSA) (Howarth and Ting, 2008). When ^AP^SSTR3^GFP^ under the control of the EF1α promoter was stably expressed in IMCD3 cells by single integration at the FlpIn locus, agonist-dependent exit of SSTR3 from cilia was undetectable. Molecular counting of GFP and comparison of immunofluorescence intensities revealed that pEF1α-driven expression resulted in SSTR3 levels that were an order of magnitude greater than in neurons (Fig. **1C**, top). Congruently, pEF1α-driven ^AP^SSTR3^GFP^ expression resulted in a near doubling of cilia length, likely due to protein overload driving ciliary membrane expansion and compensatory axoneme growth (Fig. **1C**, bottom) (Guadiana et al., 2013). To express SSTR3 at levels closer to those found in neurons, we tested a variety of weak promoters and found that an EF1α promoter lacking the TATA box (pEF1α^Δ^) produced ciliary amounts of SSTR3 similar to those found in neurons (Fig. **1C, top**). pEF1α^Δ^-driven expression of SSTR3 did not alter ciliary length (Fig. **1C**, bottom). IMCD3-[pEF1α^Δ^-^AP^SSTR3^NG^] cells recapitulated SSTR3 exit from cilia upon sst addition with nearly identical kinetics as in hippocampal neurons (Fig. **1B** and **S1D**). Similarly, the Hedgehog signaling-dependent exit of GPR161 was recapitulated by expressing ^AP^GPR161^NG3^ from the δ-crystallin promoter (Fig. **1D**). pEF1α^Δ^-driven expression of NPY2R^NG^ and MCHR1^NG^ yielded low ciliary levels (Fig. **S1E**) with similar exit kinetics (Nager et al., 2017), thus demonstrating the broad applicability of low-expression promoters for studying the dynamics of ciliary GPCRs.

Calibration of the NeonGreen signal with recombinant proteins spotted on glass slides (Fig. **1E**) allowed the measurement of absolute levels of GPCRs per cilia (Fig. **S1F-H**) and, together with pulse-chase labelling with mSA-647, enabled a specific quantitation of signal-dependent exit rates at close to 500 molecules per hour (Fig. **1F-G**, Movie **S1**). In support of the precision of our absolute quantitation, using pEF1α^Δ^-driven SSTR3^NG^ or SSTR3^GFP^ and independent calibrators yielded very similar numbers of SSTR3 molecules per cilium (Fig. **1C**).

Further highlighting the power of AP and NG-tagged GPCRs, the increased signal-to-noise ratio afforded by the direct labeling with NG or mSA647 compared to immunofluorescence (Fig. **S1I**) made it possible to detect very low abundance proteins whose presence in cilia escapes detection by traditional immunostaining techniques (Fig. **S1J**). The decreased threshold of detection when pulse-labeling with mSA647 ensures a more faithful visualization of exit kinetics than when exit is monitored by immunostaining (Fig. **1B and F**).

In the absence of agonist, the NG fluorescence of ^AP^SSTR3^NG^ increased over time whereas the signal from pulse labeling with mSA647 remained constant for 6h (Fig. **S1K**). In the presence of agonist, the exit kinetics of ^AP^SSTR3^NG^ were slower when monitored by direct visualization of the NG tag than by pulse-labeling with mSA647 (Fig. **S1K**). Since pulse labeling only reports on SSTR3 exit while the NG signal measures the total ciliary levels, these results indicate that the newly synthesized ^AP^SSTR3^NG^ continues to enter cilia during the course of the experiment. We surmise that our previous attempts to assay signal-dependent exit of SSTR3 using strong promoters failed because the entry of newly-synthesized GPCRs outpaced the slow exit kinetics.

### Sorting complexes for ciliary entry and exit

The low expression systems made it possible to validate the sorting complexes that carry out ciliary entry and exit. IFT-A is a complex of 6 protein with structural elements that suggest a common ancestry with coat complexes (Jékely and Arendt, 2006; van Dam et al., 2013) and IFT-A is recruited to membranes by the PI(4,5)P2-binding protein Tulp3 (Mukhopadhyay et al., 2010). Although IFT-A is often described as the central mediator of retrograde transport (Lechtreck, 2015), IFT-A and TULP3 are required for the import of many GPCRs into cilia (Mukhopadhyay et al., 2010, 2013; Loktev and Jackson, 2013; Fu et al., 2016; Hwang et al., 2017). We confirmed that Tulp3 was required for ciliary entry of SSTR3, NPY2R and MCHR1 (Fig. **2A** and **S2A-C**) and refined indirect interaction data by showing that the ciliary targeting signal of SSTR3 encoded within the third intracellular loop (i3) (Berbari et al., 2008a) is specifically and directly recognized by the purified IFT-A complex (Fig. **2B**) or by the IFT-A subunit IFT140 overexpressed in HEK cells (Fig. **S2D**). We conclude that IFT-A/Tulp3 functions as a coat adaptor complex that mediates GPCR entry into cilia by directly recognizing sorting signals (Fig. **2C**).

**Figure 2.**
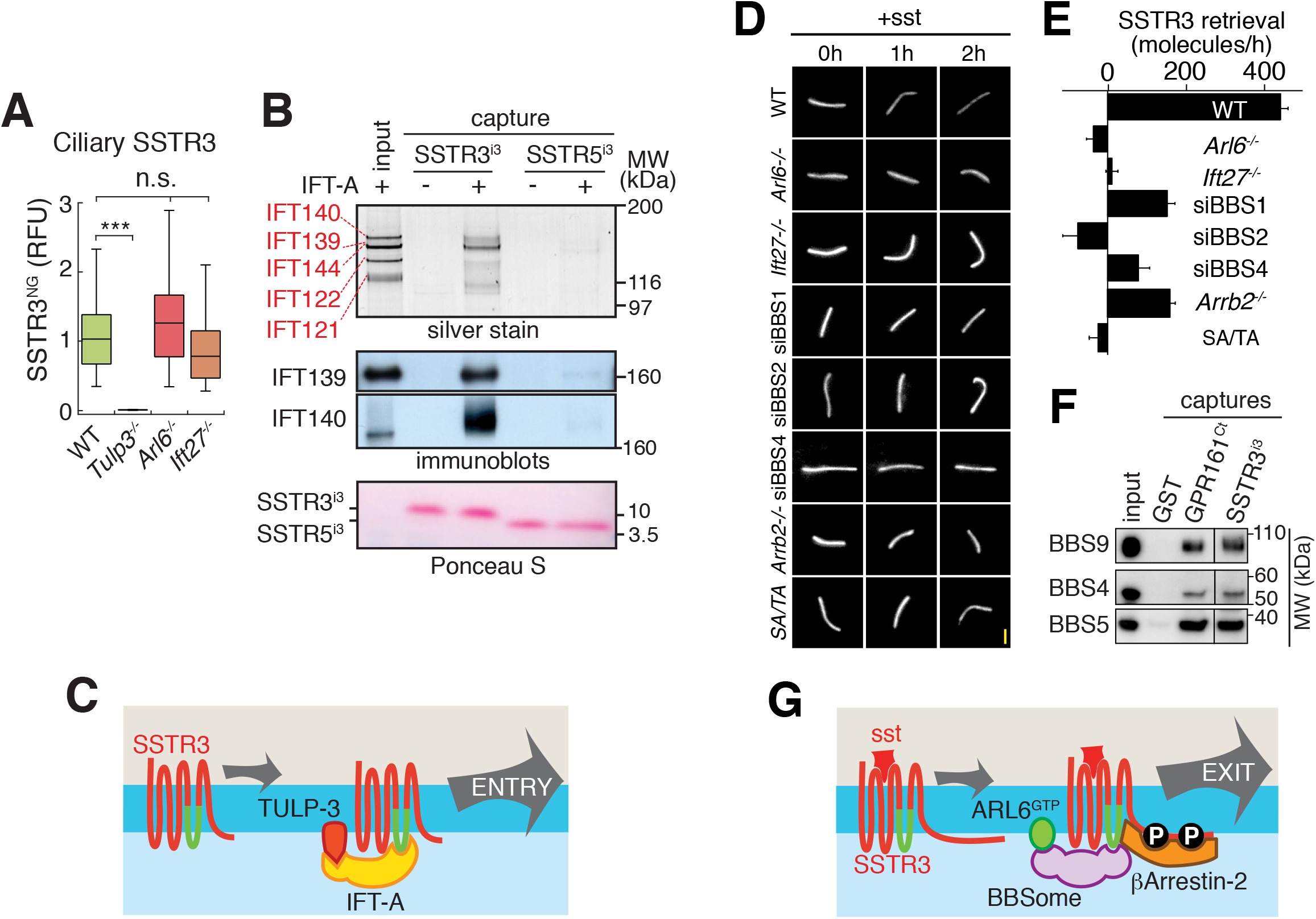
Roles of IFT-A and BBSome in ciliary entry and exit (**A**) Tulp3 is required for ciliary entry of SSTR3. Box plots of ciliary ^AP^SSTR3^NG^ intensities measured by NG fluorescence (Relative Fluorescent Unit, RFU) for various IMCD3 lines. ***Tulpĩ^/-^*** cells were fixed to identify cilia using anti-acetylated tubulin staining, all other cells were imaged live as cilia were readily identified in the NG channel. NG fluorescence was not affected by fixation (Fig. **S2B**). Asterisks indicate ANOVA significance values; *** ***P*** < 10^−4^, n.s. ***P*** > 0.05. ***N*** = 18-59 cilia. (**B**) IFT-A directly recognizes SSTR3^i3^. The IFT-A complex was purified from IMCD3-[LAP-IFT43] cells and incubated with beads coated with GST-SSTR3^i3^ or GST-SSTR5^i3^. Captured materials were eluted by cleaving off the beads and visualized by silver stain and immunoblotting. 5 input equivalents were loaded in the eluate lanes. (**C**) (**C**) Model of ciliary entry. (**D-E**) BBSome subunits were depleted by siRNA, ***Arl6, Ift27***and ***β-arrestin 2 (Arrb2)*** genes were knocked out by genome editing and SA/TA denotes a phosphomutant of the C-tail of SSTR3 that is unable to bind to β-arrestin 2. (**D**) Representative time series of ^AP^SSTR3 pulse-labeled with mSA647 under different conditions. Scale bar: 2 μm (**E**) Absolute retrieval rates were calculated by linear fitting of retrieval kinetics measured form SSTR3 pulse-chase labeling as in (**D**) (see Methods and (Nager et al., 2017)). Error bars: error of the fit. ***N***=10-35 cilia. (**F**) BBSome purified to near-homogeneity from bovine retina was incubated with glutathione beads coated with GST, GST-GPR161^Ct^ and GST-SSTR3^i3^. Captured materials were cleavage-eluted and immunoblotted. 3 input equivalents were loaded in the eluate lanes. (**G**) Signal-dependent retrieval requires the joint activities of Arl6-GTP, BBSome and -arrestin 2.

Consistent with the requirement for the GPCR activation sensor β-arrestin 2 in GPR161 retrieval (Pal et al., 2016), signal-dependent retrieval of SSTR3 required β-arrestin 2 (Fig. **2D-E** and **S2E-F**) (Green et al., 2016).

The BBSome, a complex of eight Bardet-Biedl Syndrome (BBS) proteins, resembles coat adaptors at the structural level and polymerizes into a planar coat upon recruitment to membranes by the GTP-bound form of the small GTPase Arl6/BBS3 (Jin et al., 2010). The function of the BBSome in entry vs. exit remains controversial. While BBSome mutants have decreased ciliary levels of the GPCRs somatostatin receptor 3 (SSTR3), melanin concentrating hormone receptor 1 (MCHR1), and NPY2R and of the polycystic kidney disease protein PKD1 (Berbari et al., 2008b; Loktev and Jackson, 2013; Su et al., 2014), GPR161, SMO and D1R fail to exit cilia in BBSome or Arl6 mutants (Zhang et al., 2011; Liew et al., 2014; Eguether et al., 2014; Yee et al., 2015; Nager et al., 2017). Finally, systematic studies find that some proteins accumulate while others are depleted from *Bbs* mutant cilia (Mick et al., 2015; Lechtreck et al., 2013; Datta et al., 2015). In our near-endogenous expression systems, deletion of Arl6 or of the candidate Arl6 activator Ift27/BBS19 or of β-arrestin2 did not reduce the steady-state ciliary levels of SSTR3 (Fig. **2A** and **S2F**). Instead, the BBSome, Arl6 and Ift27 were required for the signal-dependent retrieval of SSTR3 and GPR161 (Fig. **2D-E** and **S2G-H**). The carboxy-terminal tail of GPR161 (GPR161^Ct^) and the third intracellular loop of SSTR3 (SSTR3^i3^) directly interacted with purified BBSome (Fig. **2F**) (Jin et al., 2010), suggesting that BBSome coats sort signaling receptors through the direct recognition of cytoplasmic determinants. Since SSTR3 and GPR161 are recognized by β-arrestin 2 in a signal-dependent manner (Pal et al., 2016; Roth et al., 1997), we conclude that the signal-dependent retrieval of GPR161 and SSTR3 is jointly and directly mediated by β-arrestin 2 and the BBSome (Fig. **2G**).

### Signal-dependent BBSome redistribution to the tip of cilia triggers GPCR retrieval

To characterize the mechanisms of transition zone crossing by exiting GPCRs, we first sought to determine how BBSome coats facilitate signal-dependent retrieval. The movement of trains consisting of the intraflagellar transport complex B (IFT-B) can be visualized by imaging foci of the IFT-B subunit IFT88 tagged with NeonGreen traveling in the anterograde and retrograde direction inside cilia (Movie **S2**). Imaging of nematode, *Chlamydomonas* and mammalian cilia has shown that BBSome foci frequently co-move with IFT-B foci (Ou et al., 2005; Lechtreck et al., 2009; Liew et al., 2014; Williams et al., 2014), suggesting coupling between the two complexes. To follow the dynamics of the BBSome and of IFT-B during SSTR3 and GPR161 signal-dependent retrieval, we expressed ^NG3^BBS5 or ^NG3^IFT88 at near-endogenous levels (Fig. **3A** and **S2I**, Movie **S3**). In the absence of signaling, the BBSome and the IFT-B complex localized in a punctate pattern along the cilium (Fig. **3B-C**, **S2J-L**). Unexpectedly, SSTR3 activation led to a 4-fold enrichment of BBSome and a 2-fold enrichment of IFT-B at the tip (Fig. **3B-C**, **S2J-L**). Likewise, activation of the Hedgehog pathway resulted in BBSome accumulation at the ciliary tip (Fig. **3C** and Movie **S4**). Similar to ^NG3^BBS5, endogenous BBS9 became enriched at the tip upon activation of the Hedgehog pathway or of SSTR3 (Fig. **3D**). The BBSome thus joins a select group of Hedgehog factors that localize to the tip in a signal-dependent manner, consisting of Gli2, Gli3, SuFu and Kif7.

**Figure 3.**
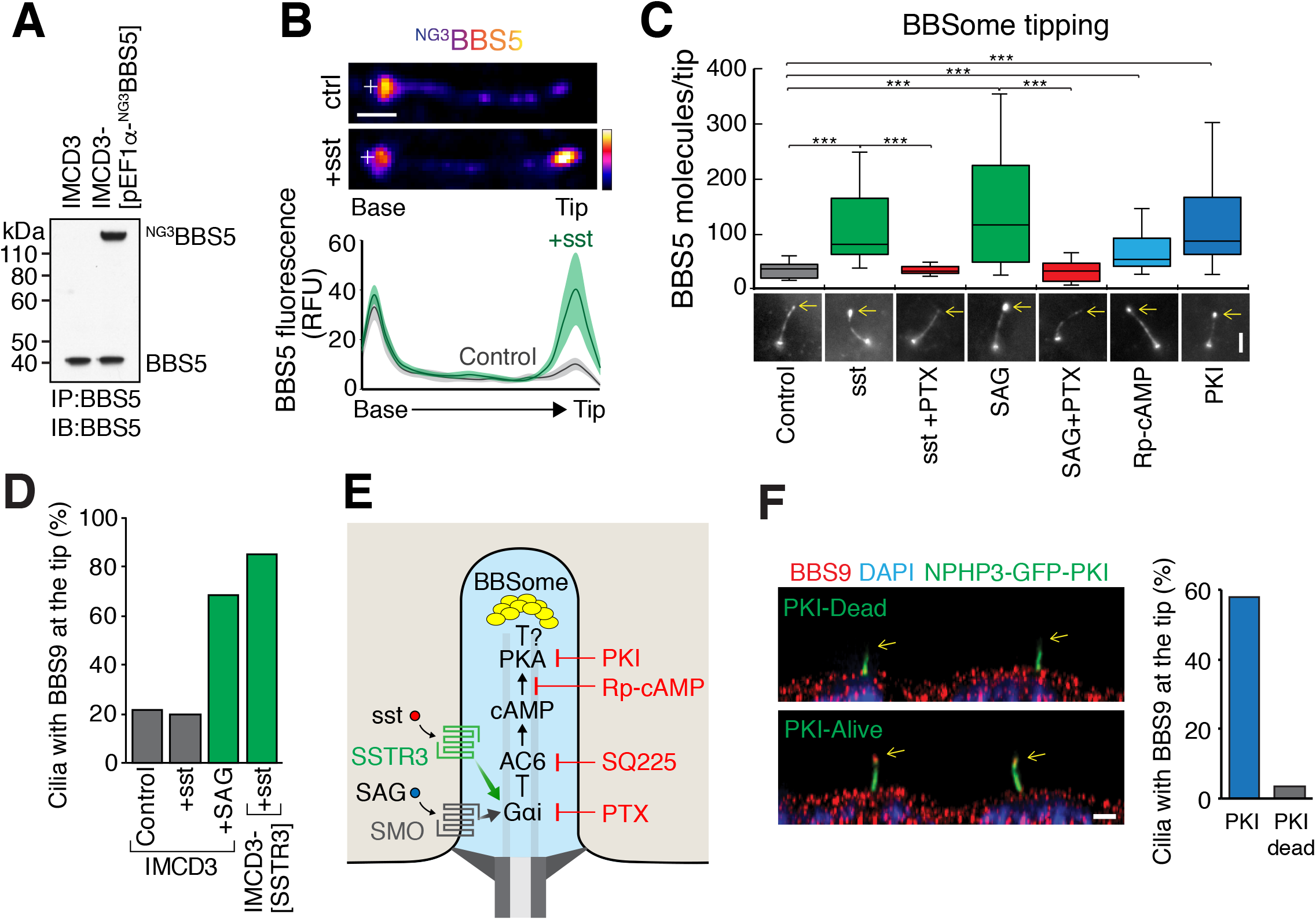
A Gαi-PKA axis promotes BBSome tip accumulation (**A**) Near endogenous expression of ^NG3^BBS5. IMCD3 and IMCD3-[pEF1α-^NG3^BBS5] cells were subjected to immunoprecipitation with an anti-BBS5 antibody and lysates and eluates were immunoblotted for BBS5. Molecular weights (kDa) are indicated on the right. Measurement of band intensities with Image Lab (Bio-Rad) indicates that the molar ratios between ^NG3^BBS5 and endogenous BBS5 is 1.27. (**B**) IMCD3-[pEF1α^Δ^-^AP^SSTR3, pEF1α-^NG3^BBS5] were treated with sst or vehicle for 40 min. Top: Representative images of cilia from live cells. ^NG3^BBS5 is in fire scale and a white cross marks the location of the basal body (see S2J). Bottom: linescans of ^NG3^BBS5 fluorescence intensities along cilia of live cells. The line marks the average intensity along length-normalized cilia. The shaded area shows the 95% confidence interval. ***N*** = 20-29 cilia. Scale bar: 1 μm. (**C**) ^NG3^BBS5 tip fluorescence intensities were quantified in live cells after 40 min of incubation with sst, SAG, Rp-cAMPs or PKI. Cells were preincubated with Pertussis toxin (PTX) for 16 h to fully inactivate G i. Representative images are shown in the lower panels with the tip marked by a yellow arrow. The total number of BBS5 molecules at the tip was calculated using the NG3 calibrator and the measured ratio of ^NG3^BBS5 to total BBS5. The whiskers represent 1.5x the interquartile range. Asterisks indicate Mann Whitney test significance values; *** ***P*** < 0.0005. ***N*** = 20-29 cilia from 3 independent experiments. Scale bar: 2 μm. (**D**) IMCD3 or IMCD3-[pEF1α^Δ^-^AP^SSTR3^NG^] cells were treated with vehicle, sst or SAG for 40 min before fixation. The bar graph shows the percentage of BBS9-positive tips detected by immunofluorescence staining of endogenous BBS9. Error bar: SD. ***N*** = 52-99 cilia. (**E**) The pathways downstream of Smoothened (SMO) and SSTR3 and the site of action of the pharmacological perturbations are shown. (**F**) Representative immunofluorescence images of ciliary BBS9 (red) in IMCD3-[pEF1a-NPHP3^GFP-PKI^] and IMCD3-[pEF1a-NPHP3^GFP-PKIdead^] cells (Mick et al., 2015). Optical sections were deconvolved and X-Z projections are shown. Arrows mark the tips of cilia. Percentage of BBS9 positive ciliary tip are plotted in the right panel. Scale bar: 2 μm.

To determine whether tip enrichment of the BBSome represents a necessary step in GPCR retrieval, we sought to define and manipulate the molecular mechanisms of signal-dependent tip accumulation. Since SSTR3 and Smoothened are known to couple to Gα_i_ and reduce cAMP production through Gα_i_-mediated inhibition of adenylate cyclases 5 and 6 (AC5/6) (Yasuda et al., 1992; Shen et al., 2013), we tested for the role of signaling downstream of Gα_i_ in promoting BBSome tip enrichment (Fig. **3E**). Pharmacological inhibition of Gα_i_ by Pertussis toxin (PTX) blocked SAG- and sst-induced BBSome tip accumulation (Fig. **3C**). Meanwhile, inhibition of AC6 (Fig. **S2M**) or inhibition of the cAMP-dependent protein kinase (PKA) by Rp-cAMPs or by a cell permeable PKA inhibitory peptide (myr-PKI) led to BBSome tip accumulation in the absence of GPCR activation (Fig. **3C**). Furthermore, cilia-targeted PKI (Mick et al., 2015) was sufficient to redistribute BBSome to the tip (Fig. **3F**). Together with the findings of cilia-localized AC5 and AC6 (Mick et al., 2015; Masyuk et al., 2008; Kwon et al., 2010), these results suggest that activation of Smoothened and SSTR3 reduce the tonically high levels of ciliary cAMP (Moore et al., 2016) through Gαi-mediated inhibition of AC5/6 within cilia. Since PKA was recently shown to reside and function inside cilia (Mick et al., 2015; Moore et al., 2016), we propose that PKA antagonizes the recruitment of BBSome to the tip of cilia in unstimulated cells and that activation of Gαi-coupled GPCRs promotes BBSome tip recruitment by reducing the activity of ciliary PKA (Fig. **3E**).

Importantly, pharmacological alterations of the ciliary Gα_i_-PKA axis concordantly affected signal-dependent redistribution of BBSome to the tip of cilia and signal-dependent GPCR retrieval as the rates of signal-dependent retrieval of GPR161 and SSTR3 were greatly reduced by PTX (Fig. **4A-B**) and significantly accelerated by myrPKI (Fig. **4C-D**). We note that PKA inhibition was not sufficient to trigger retrieval of GPR161 or SSTR3 in the absence of receptor stimulation (Fig. **4C-D**), suggesting the existence of mechanisms that act non-redundantly with the Gα_i_-PKA-BBSome axis (e.g. β-arrestin 2). In support of this hypothesis, SSTR3 activation was sufficient to elicit the retrieval of GPR161 with identical kinetics to Smoothened activation (Fig. **4E**). In contrast, Smoothened activation was not sufficient to promote SSTR3 retrieval (Fig. **4F**). As the mechanisms that underlie the activation of GPR161 remain unknown, it is conceivable that signaling downstream of SSTR3 (through Gα_i_ or Gβγ) triggers activation of GPR161 and that the subsequent engagement of β-arrestin 2 onto GPR161 cooperates with the Gαi-PKA-BBSome axis to promote GPR161 retrieval. In contrast, SSTR3 can only be activated by specific ligands.

**Figure 4.**
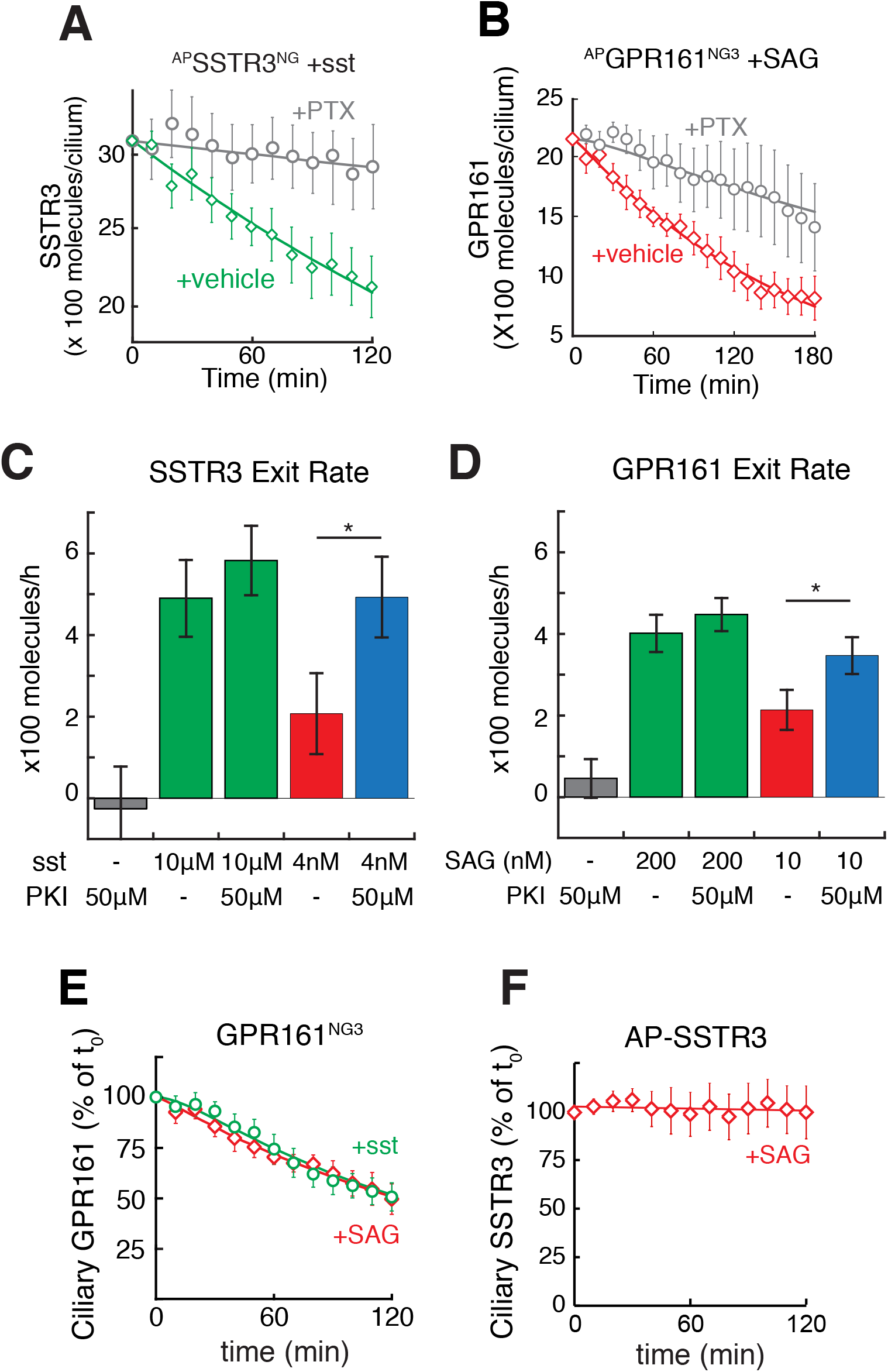
BBSome tip accumulation is required for the retrieval of SSTR3 and GPR161 (**A-B**) Pertussis toxin slows down the exit of SSTR3 (**A**) and GPR161 (**B**). IMCD3-[pEF1α^Δ^-^AP^SSTR3^NG^] or IMCD3-[pCrys-^AP^GPR161^3NG^] were pre-treated with PTX for 16 h to fully inactivate Gα¡. After agonist treatment, ciliary GPCR levels were measured in live cells as described in Fig. **1F-G**. Error bars: 95% CI. ***N*** = 10-28 cilia. (**C-D**) PKA inhibition accelerates the exit of SSTR3 (**C**) and GPR161 (**D**). IMCD3-[pEF1α^Δ^-^AP^SSTR3^NG^] or IMCD3-[pCrys-^AP^GPR161^3NG^] cells were treated for 3 h with the indicated concentrations of agonist and/or PKI. Ciliary level of GPCRs were measured by NeonGreen fluorescence before and after treatment to estimate the rate of exit. Addition of PKI together with sub-saturating concentrations of agonist significantly accelerated the GPCR exit rates to the near-maximal values observed with saturating concentrations of agonist. Error bars: Error of the fit. ***N*** = 67-112 cilia from 3 independent experiments. Asterisks indicate multiple regression significance values; *P < 0.05. (**E**) GPR161 retrieval can be triggered by sst treatment. IMCD3-[pCrys-^AP^GPR161^3NG^] cells were treated with SAG or sst for 2h. NG fluorescence was tracked in individual cilia. Data were fitted to a single exponential. Error bars: 95% CI. ***N***=10-21 cilia. (**F**) SAG treatment is not sufficient to trigger the retrieval of SSTR3. IMCD3-[pEF1α^Δ^-^AP^SSTR3^NG^] cells were treated with SAG for 2h. Stable expression of an ER-localized biotin ligase BirA enables detection of ^AP^SSTR3 by pulse-labeling live cells with Alexa647-conjugated mSA (mSA647) and tracking individual cilia. Data were fitted to a single exponential. Error bars: 95% CI. ***N*** = 12 cilia.

To further establish that BBSome tipping represents a necessary intermediate in GPCR retrieval, we sought to identify molecules that recruit the BBSome to the tip of cilia in a signal-dependent manner. The plus-end directed microtubule motor Kif7 represents a candidate tip recruitment factor because Kif7 accumulates at the tip of cilia upon Hedgehog pathway activation (Liem et al., 2009; Endoh-Yamagami et al., 2009) and is necessary and sufficient to promote tip accumulation of the Hedgehog signaling factors Gli2 and Gli3 (He et al., 2014). Furthermore, *KIF7* is a genetic modifier of Bardet-Biedl Syndrome in human patients (Putoux et al., 2011). To follow the behavior of Kif7 and the BBSome in live IMCD3 cells, we stably co-expressed ^NG3^BBS5 and Kif7 fused to the red fluorescent protein mScarlet. Smoothened activation led to the correlated co-accumulation of BBS5 and Kif7 at the tip of cilia (Fig. **5A-B**). Furthermore, in the rare instances where a second spot of Kif7 was found along cilia, possibly because part of the axoneme terminates prior to the tip, a similarly intense second spot of BBS5 was observed at the same location (Fig. **5C**). Since Kif7 depletion abolished tip accumulation of the BBSome (Fig. **5D-E**), these data suggest that Kif7 directly mediates the signal-dependent recruitment of BBSome to the tip of cilia. Alternatively, it is conceivable that structural defects in cilia of Kif7-depeleted cells indirectly affect the recruitment of BBSomes to the tip. Congruent with a Kif7-mediated recruitment of BBSome to tips of cilia, Kif7 was found to co-immunoprecipitate with several BBSome subunits (Fig. **S2N**). Given that dephosphorylation of Kif7 leads to the accumulation of Kif7 at the tip of cilia (Liu et al., 2014), we considered that phosphorylation by PKA may directly antagonizes Kif7 tip accumulation and the Kif7-BBSome interaction. Concordantly, PKI led to the correlated co-accumulation of BBS5 and Kif7 at the tip of cilia to the same extent as SAG (Fig. **5A-B**) and elevated cAMP levels decreased the Kif7-BBSome interaction (Fig. **5F**). Finally, Kif7 was required for signal-dependent retrieval of SSTR3 (Fig. **5G**). Together, these results suggest that signaling downstream of Smoothened and SSTR3 leads to Kif7 dephosphorylation and that ensuing recruitment of BBSome to the tip initiates GPCR retrieval.

**Figure 5.**
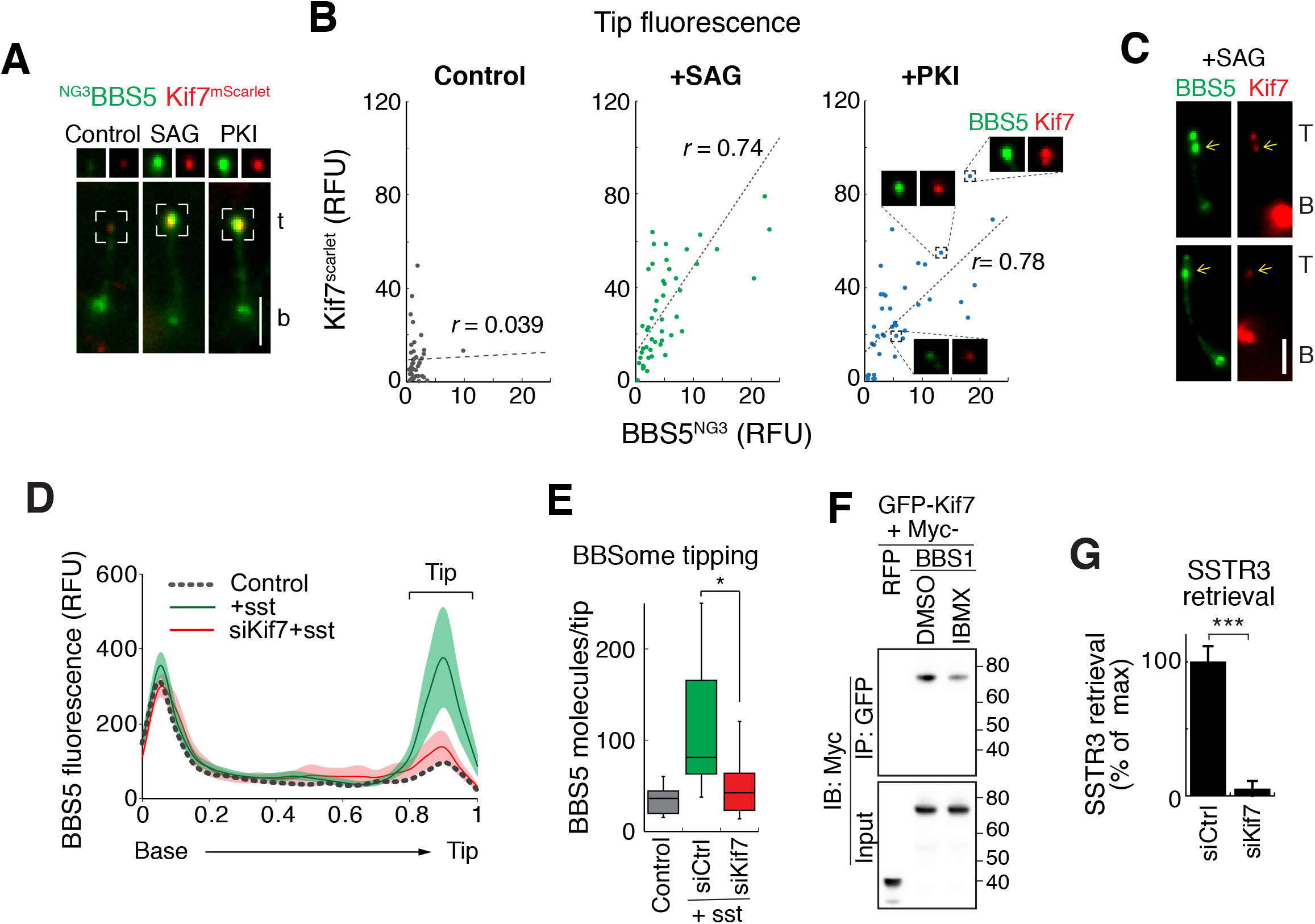
PKA antagonizes the recruitment of the BBSome to the tip of cilia by Kif7 (**A-C**) Co-recruitment of BBS5 and Kif7 to the tip of cilia in live IMCD3-[pEF1α-^NG3^BBS5; pCMV-Kif7^mScarlet^] cells. (**A**) Representative images showing SAG- and PKI-induced accumulation of ^NG3^BBS5 and Kif7^mScarlet^ at ciliary tip. Cells were treated with SAG, PKI or vehicle for 40 min before imaging. White boxes mark the ciliary tip and split channels of ^NG3^BBS5 and Kif7^mScarlet^ are shown on top. Scale bar: 2 μm. (**B**) Correlation between the fluorescence signal of ^NG3^BBS5 and Kif7^Scarlet^ at the ciliary tip. Fluorescence signals of ^NG3^BBS5 and Kif7^mScarlet^ were measured at ciliary tip of live cells after 40 min treatment with vehicle, SAG or PKI. Micrographs of ^NG3^BBS5 and Kif7^mScarlet^ at the ciliary tip for three representative data points are shown. Linear regressions (dotted lines) highlight the positive correlation between ciliary tip levels of ^NG3^BBS5 and Kif7^mScarlet^ in the presence of SAG or PKI. The Pearson correlation coefficient (r) is shown. Student’s t-test of the Pearson’s correlation coefficient (r) returned a non-significant ***P*** value under control conditions (P > 0.7) but a significant value after SAG or PKI treatment (P < 10^−5^). ***N*** = 40-49 cilia. (**C**) In SAG-treated cells where a second spot of Kif7 is occasionally found along cilia, a second spot of BBS5 was observed at the same location as Kif7. The yellow arrows mark the location of ^NG3^BBS5 foci that accumulated at the ectopic tip. Scale bar: 2 μm. (**D-E**) Kif7 is necessary for the redistribution of BBS5 to the tip of cilia. (**D**) Linescans of ^NG3^BBS5 fluorescence intensities along cilia of live cells. Cells were transfected with siRNAs for 72h and treated with sst for 40 min before live imaging of ^NG3^BBS5 fluorescence. The line marks the average intensity along length-normalized cilia. The shaded areas show the 95% confidence interval (not shown for the vehicle control). ***N*** = 20-27 cilia. (**E**) The total number of BBS5 molecules at the tip was calculated as in Fig. **3C**. Asterisks indicate Mann Whitney test significance values; * ***P*** < 0.05. ***N*** = 20-28 cilia from 3 independent experiments. (**F**) Kif7 interacts with BBSome and PKA antagonizes this interaction. HEK293 cells cotransfected with Kif7^GFP^ and ^Myc^BBS1 were treated with the cAMP phosphodiesterase inhibitor IBMX or vehicle for 30 min before lysis. Complexes were immunoprecipitated with anti-GFP antibodies. Lysates and eluates were blotted for Myc. The capture efficiency of ^Myc^BBS1 by Kif7^GFP^ was decreased 29 ± 3% upon treatment with IBMX. Molecular weights (kDa) are indicated on the right. ***N*** = 3 independent experiments. (**G**) Kif7 is necessary for SSTR3 exit from cilia. IMCD3-[^AP^SSTR3^ng^] cells were treated with siRNA targeting Kif7 or Luciferase, pulse-labeled with SA647, and imaged every 10 min following addition of sst. The resulting loss in SA647 fluorescence was plotted and linearly fitted to determine the rate of SSTR3 retrieval. Asterisks indicate Mann Whitney test significance values; * ***P*** < 0.05, *** ***P*** < 0.0005. Error bars: SD. ***N*** = 13 cilia.

### BBSome tip accumulation drives formation of cargo-laden retrograde trains

Analysis of ^NG3^BBS5 and ^NG3^IFT88 kymographs showed that anterograde BBSome trains, anterograde IFT trains and retrograde IFT trains moved processively along the length of the cilium regardless of the signaling status (Fig. **6A** and S3A-B). Yet in unstimulated cells, BBSome trains occasionally detached from retrograde IFT trains before reaching the base (Fig. **6B**) and overall, 90% of BBSome trains failed to reach the base of cilia (Fig. **6A, C** and **S3B**). Activation of Smoothened or SSTR3 or inhibition of PKA all doubled the number of BBSomes per retrograde train from 10 to 20 (Fig. **6D**) and led to a significant increase in the processivity of retrograde BBSome trains (Fig. **6C** and **S3B**). Neither the frequencies nor the velocities of BBSome trains were affected by these treatments (Fig. **6E-F**). Meanwhile, SAG increased the number of IFT-B particles per retrograde train from 62 to 78 (Fig. **6G**). The addition of 106 BBSomes and 206 IFT-B to the tip upon SSTR3 activation (Fig. **3C** and **S2L**) suggests that the signal-dependent accumulation of BBSome and IFT-B at the tip drives the growth of retrograde trains by increasing the concentration of precursors at the site of assembly. Similar to the BBSome and IFT-B, Arl6 underwent signal-dependent tip accumulation (Fig. **6H**). As Arl6 was required for IFT88 tipping but not BBSome tipping (Nager et al., 2017) (Fig. **S2L**), Arl6 may recruit IFT-B particles to the tip by increasing the affinity of the BBSome for IFT-B.

**Figure 6.**
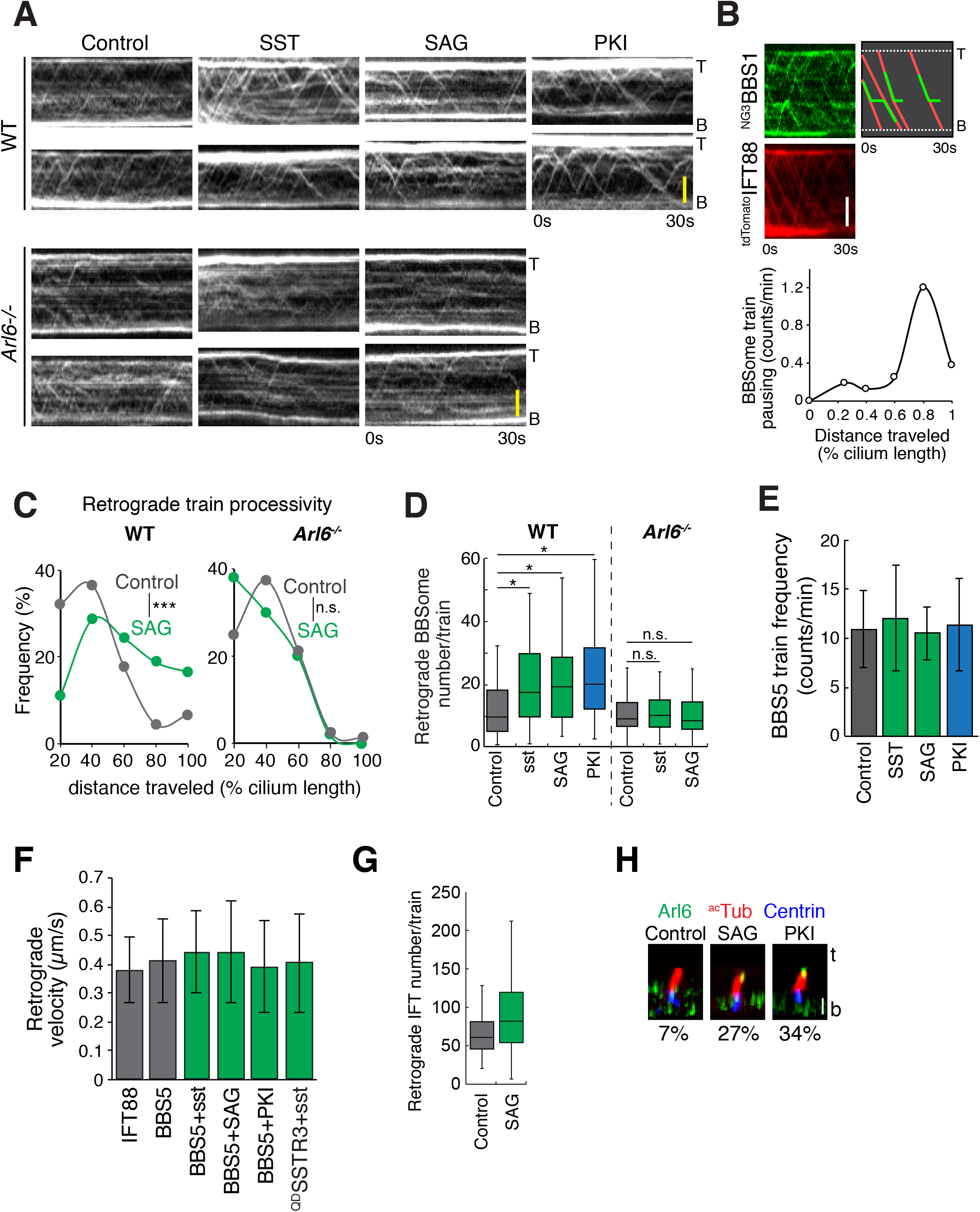
GPCR signaling and Arl6 drive assembly of large, processive retrograde BBSome trains (**A**) Representative kymographs of BBSome train movement. WT or ***Arl6^-/-^*** IMCD3-[pEF1α-^NG3^BBS5; pEF1α^Δ^-^AP^SSTR3] cells were treated with vehicle, sst, SAG or PKI for 40 min before imaging at 4 Hz for 30 s. Scale bar: 2 μm (**B**) Representative kymograph from an IMCD3-[pEF1α-^NG3^BBS1, pCMV-^tdTomato^IFT88] cell showing the co-movement and the uncoupling between IFT-B and BBSome trains in untreated cells. Nearly 20% of BBSome trains displayed a distinct pause. (**C-D**) WT or ***Arl(?^/-^*** IMCD3-[pEF1α-^NG3^BBS5; pEF1α^Δ^-^AP^SSTR3] cells were treated with vehicle, sst, SAG or PKI for 40 min before imaging. (**C**) The processivity of retrograde BBSome trains was measured by deconvolving kymographs into anterograde and retrograde components (see methods). The distance traveled by each retrograde train (normalized to the length of cilia) was estimated by manual inspection of the retrograde kymographs. ***N***= 52-91 cilia form 3 independent experiments. Asterisks indicate the significance values of Mann-Whitney Utest applied to the entire distribution; *** ***P*** < 0.0005, n.s. ***P*** > 0.05. (**D**) The fluorescence intensity of ^NG3^BBS5 retrograde trains was extracted from deconvolved kymographs and the total number of BBS5 molecules per train was calculated using the NG calibrator (see methods). The whiskers represent 1.5x the interquartile range. ***N*** = 52-91 cilia from 3 independent experiments. Asterisks indicate Mann-Whitney ***U*** test significance values; *** ***P*** < 10^−4^, n.s. ***P*** > 0.05. (**E**) Treatment with sst, SAG or PKI did not change the frequency of retrograde BBSome trains. Error bars: SD. ***N*** = 9-18 cilia. Pairwise Mann-Whitney tests fail to show significant differences between any two conditions (P > 0.1). (**F**) Retrograde velocities of IFT trains, BBSome trains and single SSTR3 molecules. IMCD3-[pEF1α-^NG3^IFT88], IMCD3-[pEF1α-^NG3^BBS5; pEF1α^Δ^-^AP^SSTR3] or IMCD3-[pEF1α^Δ^-^AP^SSTR3^NG^] were treated with vehicle, sst, SAG or PKI for 40 min before imaging. IFT and BBSome train velocities were extracted from kymographs (see methods). ^QD^SSTR3 velocities were measured from persistent retrograde movements lasting more than 6s. Error range: SD. ***N*** = 9-18 cilia. Pairwise Mann-Whitney tests fail to show significant differences between any two conditions (P > 0.1). (**G**) The number of IFT88 molecules per retrograde train was measured in IMCD3-[pEF1α-^NG3^IFT88] cells treated with vehicle or SAG. Counting of molecules is detailed in Methods. (**H**) Arl6 immunofluorescence of cells treated with vehicle, SAG or PKI. Optical sections were deconvolved and X-Z projections are shown. The percentages of Arl6-positive tips are indicated below the micrographs. ***N*** = 88-118 cilia from 4 to 5 microscopic fields.

PKI was sufficient to trigger tip accumulation of BBSome and Arl6 as well as the formation of large processive retrograde BBSome trains (Fig. **6D, H** and **S3B**), suggesting that recruitment of Arl6 and BBSome to the tip may be sufficient to initiate the assembly of large processive retrograde IFT/BBSome trains. Because Arl6 and its candidate activator Ift27 were required for the signal-dependent formation of large processive BBSome trains (Fig. **6C-D** and **S3C**), we propose that BBSome coats polymerized upon Arl6-GTP binding become stably coupled to IFT-B trains to generate the large, signal-dependent, and processive retrograde BBSome/IFT trains.

Tip redistribution upon GPCR activation was not limited to the BBSome and IFT-B as GPR161 (Fig. **7A-B**) also underwent tip redistribution. Photobleaching the cilium exclusive of the tip revealed an enrichment of GPR161 at the tip upon Hh pathway activation (Fig. **7A-B**). Recovery kinetics show that GPR161 at the tip became less dynamic after SAG treatment (Fig. **7C**). In support of an association between BBSome and activated GPCRs at the tip of cilia, we find that WGA-mediated immobilization of membrane proteins increases the amount of BBSome at the tip, most likely because BBSome-GPCR complexes are unable to leave the tip (Fig. **7D**). Furthermore, WGA treatment increases the amount of BBSome throughout the length of cilia (Fig. **7D**), indicating that BBSome trains become trapped by immobilized GPCRs.

**Figure 7.**
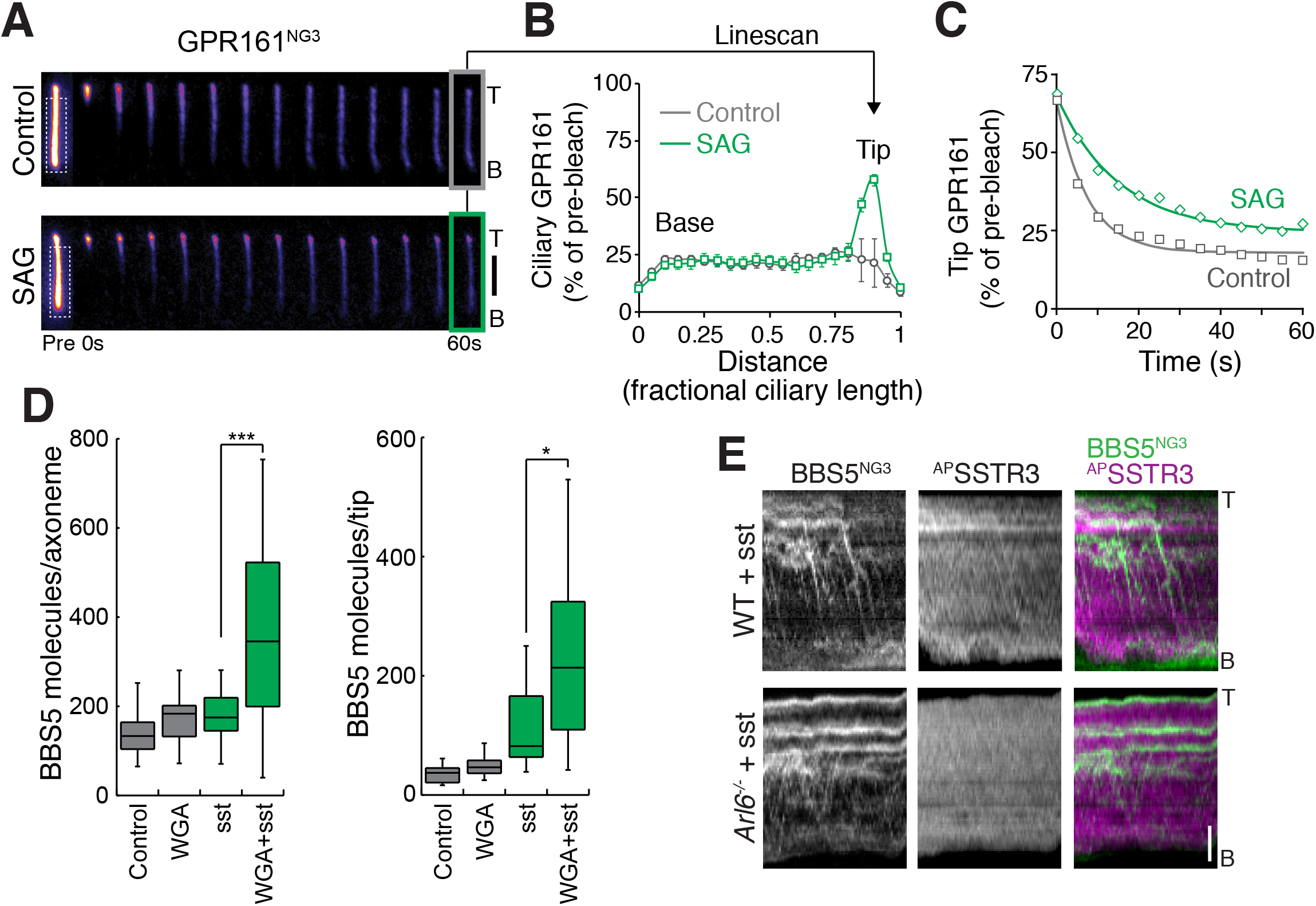
Signaling promotes coupling between BBSome and cargoes (**A**) Representative images showing ciliary GPR161^3NG^ fluorescence recovery after photobleaching (FRAP). IMCD3-[pCrys-^AP^GPR161^3NG^] cells were treated with vehicle or SAG for 40 min before imaging. A white dotted box in the first image indicates the photobleaching area that covers more than 80% of the cilia except the tip. After photobleaching, ciliary GPR161^3NG^ images are acquired every 5 s. Scale bar: 2 μm (**B**) Linescans of GPR161^3NG^ fluorescence intensities along cilia at 60 s after photobleaching. The grey and green lines mark the average intensities along length-normalized cilia for control and SAG treated cells respectively. Error bar: SD. ***N*** = 10-11 cilia. (**C**) GPR161^3NG^ fluorescence at the tip was measured and the decay of fluorescence signal over time in control and SAG-treated cells was plotted. Data were fitted to a single exponential to calculate the half-life. t_?/2_[control] = 5.14 +/-0.95 s, t_?/2_[SAG]= 10.47 +/-1.76 s. Error range: error of fit. ***N*** = 10-11 cilia. (**D**) ^NG3^BBS5 fluorescence was measured in live cells after 40 min of incubation with WGA, sst, or both. The total number of BBS5 at the tip (right) or axoneme (left) was calculated using the NG3 calibrator and the measured ratio of ^NG3^BBS5 to total BBS5. Asterisks indicate Mann Whitney test significance values; * ***P*** < 0.05, *** ***P*** < 0.0005. ***N***= 20-25 cilia from 3 independent experiments. (**E**) Representative kymographs from mSA647-labeled WT or ***Arl6^-/-^*** IMCD3-[pEF1α-^NG3^BBS5; pEF1α^Δ^-^AP^SSTR3] cells treated with sst for 1 h before imaging. The cells stably expressed an ER-localized biotin ligase BirA to enable visualization of ^AP^SSTR3 by mSA647 labeling.

The bright and processive retrograde BBSome tracks observed upon sst addition frequently overlapped with faint tracks of SSTR3 (Fig. **7E**, Movie **S5**). No co-movement was observed between the faint BBSome trains and SSTR3 in the absence of Arl6 (Fig. **7E** and **S3D**). These observations suggest that BBSome/Arl6 coats capture cargoes and move them from tip to base of cilia upon coupling to retrograde IFT trains.

### Signaling promotes the processive retrograde movement of GPCRs

In bulk fluorescence imaging, the few fluorescent GPCRs that are being trafficked tend to be obscured by the fluorescent GPCRs that remain inside cilia. We measured that the largest retrograde trains contain close to 50 BBSomes. Assuming a 1:1 stoichiometry between BBSome and cargo, this indicates that the brightest retrograde cargo tracks can carry at most 50/3129=1.6% of the total ciliary SSTR3. It is therefore expected that such tracks will be extremely faint by bulk imaging (Fig. **7E**). To overcome the limitations inherent to ensemble imaging, we set out to visualize the molecular events that underlie GPCR retrieval at singlemolecule resolution. Combining site-specific biotinylation of AP-tagged GPCRs (Ye et al., 2013) with streptavidin-coupled quantum dots (^SA^Qdot) enabled imaging at 2 Hz for over 20 min (compared to 30-60s when using mSA647) (Fig. **S3E**). By blocking most surface-exposed biotin groups with mSA before labeling with ^SA^Qdot, the vast majority of cilia bore no ^SA^Qdot and cilia bearing one ^SA^Qdot are expected to possess a single biotinylated ^AP^GPCR (Fig. **8A**). Tracking of ^SA^Qdot-labeled GPCRs (^QD^GPCRs) demonstrated that their dynamics were unperturbed by ^SA^Qdot labeling, consistent with a single valence of ^SA^Qdot per GPCR.

**Figure 8.**
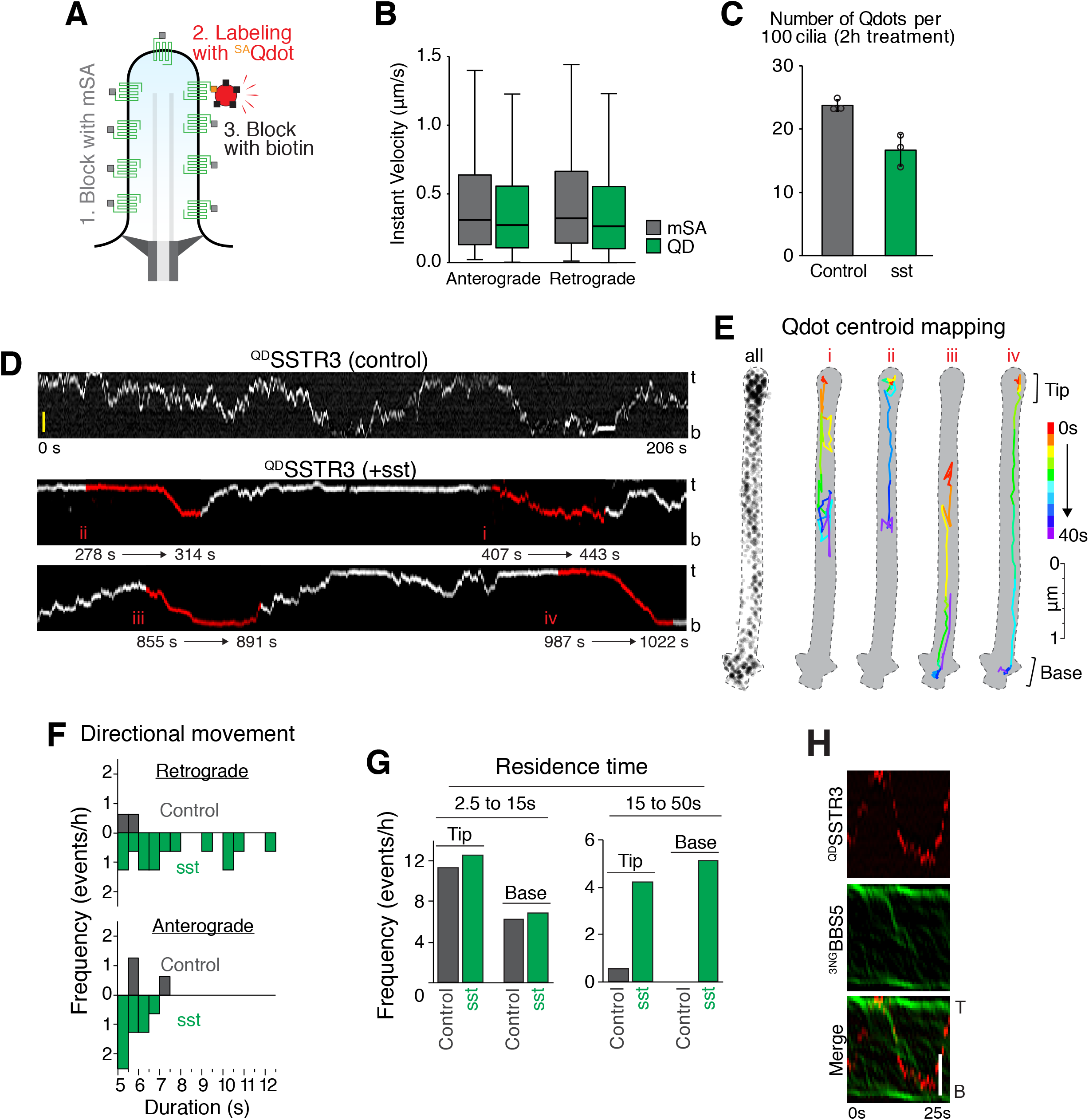
Activated GPCRs undergo processive retrograde movements, and confinements at base and tip (**A**) Diagram of the Qdot labeling strategy. IMCD3-[pEF1α^Δ^-^AP^SSTR3^NG^] cells stably expressing BirA-ER were first treated with unlabeled mSA to passivate the surface-exposed biotinylated ^AP^SSTR3. ^SA^Qdots were then added to the medium to label the GPCRs newly arrived at the surface. Finally, biotin was added to the medium to passivate the excess SA on Qdots. (**B**) The diffusive properties of SSTR3 are not altered by Qdot labeling. The instantaneous velocities of mSA647-labeled ^AP^SSTR3^NG^ and Qdot655-labeled ^AP^SSTR3^NG^ were measured by single molecule tracking in the absence of sst. Error bars: SEM. (**C**) Qdot labeling does not alter the exit rate of SSTR3. IMCD3-[pEF1α-^AP^SSTR3^GFP^] cells were sparsely labeled with Qdot655 as described in Methods and treated with vehicle or sst for 2h before fixation. The number of Qdot per cilium was counted in both vehicle or sst treated condition. ***N*** = 263-303 cilia from three independent experiments. Error bars: SEM. (**D**) Representative kymographs showing the movements of ^SA^Qdot-labeled ciliary ^AP^SSTR3^NG^ (^QD^SSTR3) in vehicle-or sst-treated cells. Red labels and line coloring highlight four characteristic movement behaviors. Scale bar: 2 μm. (**E**) Centroid mapping of ^QD^SSTR3. Left: the contour of the cilium was traced as a dotted line that captures all ^qd^SSTR3 positions. (i) diffusive movement, (ii) tip confinement followed by retrograde movement, (iii) retrograde movement followed by base confinement and return into the cilium, and (iv) tip confinement followed by fully processive retrograde movement and base confinement. The time dimension is color-coded from red to purple. (**F-G**) Signaling increases tip confinement and processive retrograde movement of SSTR3. Cells were treated with sst (green, sst) or vehicle (grey, Control) for 40 min before imaging was initiated for 10 to 20 min. (**F**) Durations of persistent movement events for ^QD^SSTR3 in anterograde and retrograde directions. (**G**) The durations of confinement events for ^QD^SSTR3 at ciliary tip or base were binned into two categories. ***N*** =12 cilia for each condition. (**H**) Representative kymographs showing the co-movement between a single ^QD^SSTR3 and a BBSome retrograde train. IMCD3-[pEF1α-^NG3^BBS5; pEF1α^Δ^-^AP^SSTR3] cells were treated with sst for 40 min before imaging.

First, consistent with prior single-molecule tracking studies (Ye et al., 2013; Milenkovic et al., 2015), ^QD^SSTR3 displayed a diffusive behavior in the absence of agonist (Fig. **8D**, **S4A** and Movie **S6**) and instant velocities of diffusing GPCRs were nearly identical when labeled by mSA647 or ^SA^QD655 (Fig. **8B**). Second, congruency of exit rates between bulk imaging and ^QD^SSTR3 imaging indicated that ^SA^Qdot labeling did not impair exit (Fig. **8C**). Mapping the centroid of the Qdot enabled a 7 nm measurement precision (Fig. **S3F**) that resolved the lateral displacement of ^QD^SSTR3 around the 250 nm diameter of the ciliary membrane during diffusive events (Fig. **8Ei**). Addition of sst led to ^QD^SSTR3 undergoing frequent confinements at the base and tip as well as processive retrograde movements (Fig. **8D-E, ii-iv** and **S4B** and Movie **S7**). Although we occasionally observed apparent directional movement during brief periods of time for ^QD^SSTR3 in control-treated cells (Fig. **8D** and **S4A**), persistent retrograde movements of ^QD^SSTR3 lasting 6 s or more were strictly dependent on SSTR3 activation (Fig. **8F** and **S3G**), confirming that signaling drives formation of cargo-laden IFT/BBSome retrograde trains. This signal-dependent increase in long processive movements was not observed for anterograde movements (Fig. **8F**). Processive transport events did not exhibit lateral displacement (Fig. **8Eii-iv**), strongly suggesting that each cargo-laden IFT/BBSome train moves along a single axonemal microtubule. GPCRs occasionally resided for several seconds at either the tip or the base, and the frequency of confinement events −defined as residence exceeding 15s– significantly increased in the presence of sst (Fig. **8G** and **S3H**). Since processive retrograde transport events were frequently preceded by tip confinement (Fig. **8Eii** and **iv**), it is likely that tip confinement of GPCRs reflects a step of cargo capture by BBSome coats assembling at the tip. Nevertheless, ^QD^SSTR3 exhibited retrograde movements starting at any point along the cilium (Fig. **8Eiii** and **S4B**), indicative of activated GPCRs hopping onto retrograde BBSome trains.

Similar to ^QD^SSTR3, ^QD^GPR161 underwent mostly diffusive behavior in unstimulated cells, and addition of SAG led to frequent retrograde processive transport events (Fig. **S4C-D**). Consistent with IFT/BBSome trains powering the retrograde transport of activated GPCRs, long processive retrograde movements of ^QD^GPR161 were absent when *Arl6* was deleted (Fig. **S4E**). The average velocity of the processive retrograde movements of ^QD^SSTR3 was similar to the velocities of IFT and BBSome trains (Fig. **6F**), suggesting that signaling promotes coupling between cargoes and retrograde IFT trains. In support of this hypothesis, co-imaging of ^3NG^BBS5 and ^QD^SSTR3 uncovers instances of co-movement between retrograde BBSome train and ^QD^SSTR3 (Fig. **8H**). We conclude that activated GPR161 and SSTR3 are transported in the retrograde direction by the large, processive BBSome trains that couple to retrograde IFT trains upon GPCR activation.

### Arl6 and signaling enable transition zone crossing by GPR161

The expression of ^AP^GPCRs at near-endogenous levels enabled the ^SA^Qdot-mediated visualization of exit from the ciliary compartment at single-molecule resolution. In combined 21 hours of single molecule imaging, three exit events of ^QD^GPR161 were observed. In all three events, ^QD^GPR161 exit followed a stereotypical sequence of processive retrograde transport, base confinement and diffusion away from the ciliary compartment (Fig. **9A-B**, event **iii**).

**Figure 9.**
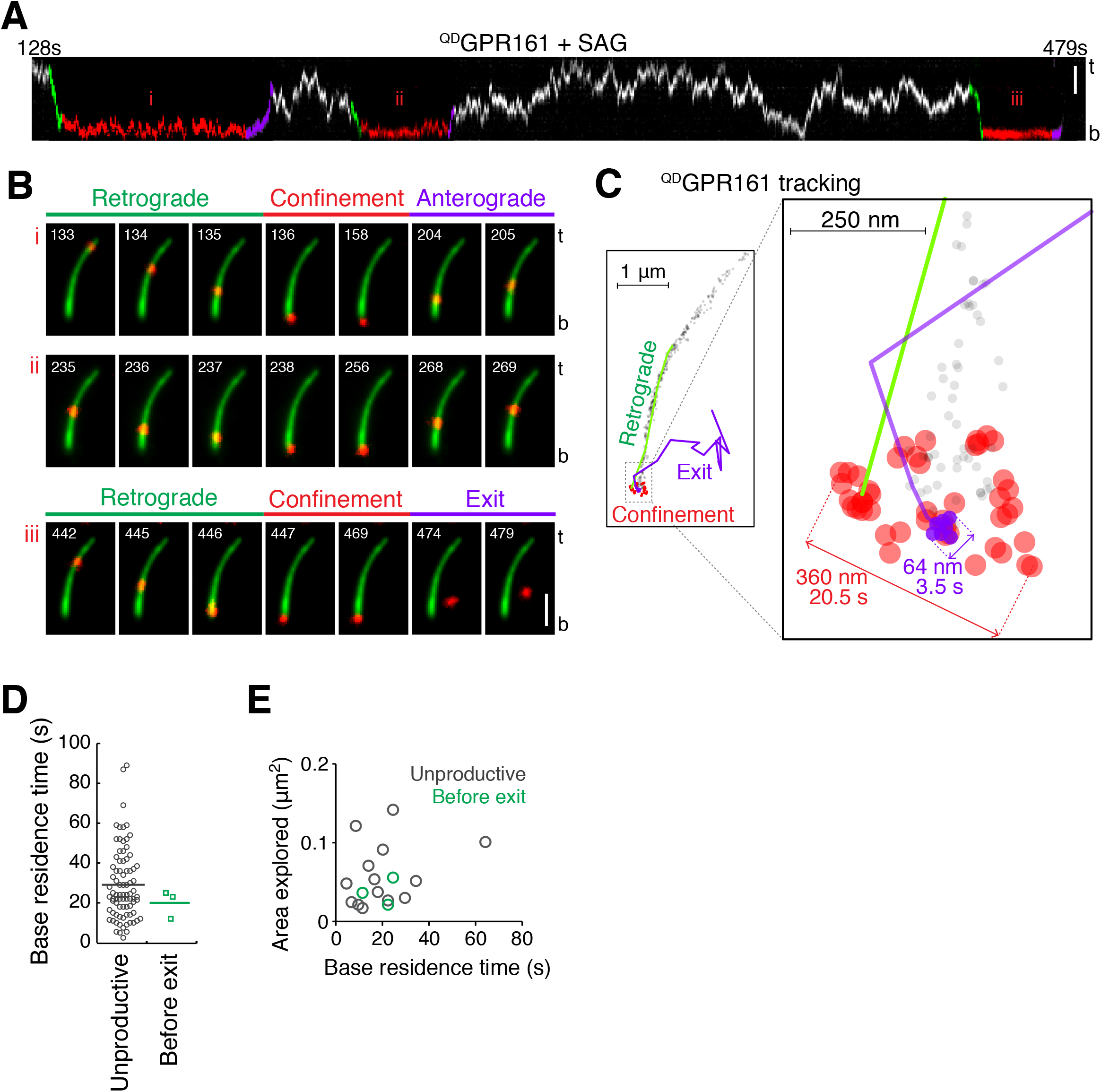
GPCR base confinement results in either exit or re-entry (**A**) Cilium kymograph of ^QD^GPR161. Cells were treated with SAG for 40 min before the start of imaging. Green, red and purple labels and line coloring indicate retrograde movements, confinement and anterograde transport (events i and ii) or exit from cilia (iii). Scale bar: 2 μm. (**B**) Time series of the three confinement events. A reference image of bulk NG fluorescence from ^AP^GPR161^NG3^ was overlaid with images of ^QD^GPR161. Time stamp (s) is in the upper left corner. Scale bar: 2 μm. t: tip. b: base. (**C**) Centroid mapping of ^QD^GPR161 during event iii. The ^QD^GPR161 locations captured during the entire imaging session are shown as grey dots. The green track represents a processive retrograde transport event that precedes confinement of ^QD^GPR161 (shown in red dots) at the base. Immediately before diffusion away from the cilium (purple track), ^QD^GPR161 was nearly immobile for 4.5 s (purple dots). (**D**) Dot plot showing the base residence time of ^QD^GPR161 during unproductive events (grey circles) or before exit (green boxes). IMCD3-[pCrys-^AP^GPR161^3NG^] cells were treated with SAG for 40 min before imaging. (**E**) Scatter plot of areas explored by ^QD^GPR161 during base residence events versus time. Grey circles represent unproductive events and green circles represent pre-exit base residence events. Linear regression shows no obvious correlation between these two variables (Pearson correlation coefficient ***r***= 0.3).

Surprisingly, many ^QD^GPCRs that underwent retrograde transport and base confinement returned into the shaft of the cilium, seemingly by processive anterograde transport (Fig. **9A-B**, events **i** and **ii**, and **8D-E** event **iii**). During two out of the three pre-exit confinement events we observed, ^QD^GPR161 first diffused within an area of less than 360 nm diameter before near-complete immobilization prior to exit (Fig. **9C**, **S5A** and Movie **S8-9**). We did not observe a clear correlation between successful exit and either base residence time (Fig. **9D**) or area explored before exit (Fig. **9E**) suggesting that completion of exit is a stochastic process. Because ^QD^GPR161 diffused within the 0.4 μm-deep focal plane after exit from the ciliary compartment, it is most likely that the GPCR moved into the plasma membrane after exit (Fig. **9C**, **S5A** and Movie **S8-9**). Exit event were observed at the expected frequency based on measurements of bulk exit rates (Fig. **S5B**), this confirms that ^SA^Qdot labeling did not interfere with exit.

Unexpectedly, the position of ^QD^GPR161 during signal-dependent base confinement events was often separate from the bulk fluorescence of GPR161^NG3^ (Fig. **10A**). Using the profile of the NeonGreen channel as a common reference (Fig. **10A-C** and **S5C**), we mapped the location of the most base-proximal position explored by ^QD^GPR161 relative to the transition zone marker Cep290 and the transition fiber marker Cep164 and found that activated ^QD^GPR161 moves into the 100 nm space between these two markers (Fig. **10A-C**). We termed the region between the 10^th^ and 50^th^ percentile of GPR161^NG3^ fluorescence the intermediate compartment as it defined the area visited by ^QD^GPR161 before exit was completed. Systematic analysis of ^QD^GPR161 position demonstrated that GPR161 enters the intermediate compartment in a signal- and Arl6-dependent manner (Fig. **10C**), thereby demonstrating that large, processive BBSome trains ferry GPR161 through the transition zone and deliver it to the intermediate compartment.

**Figure 10.**
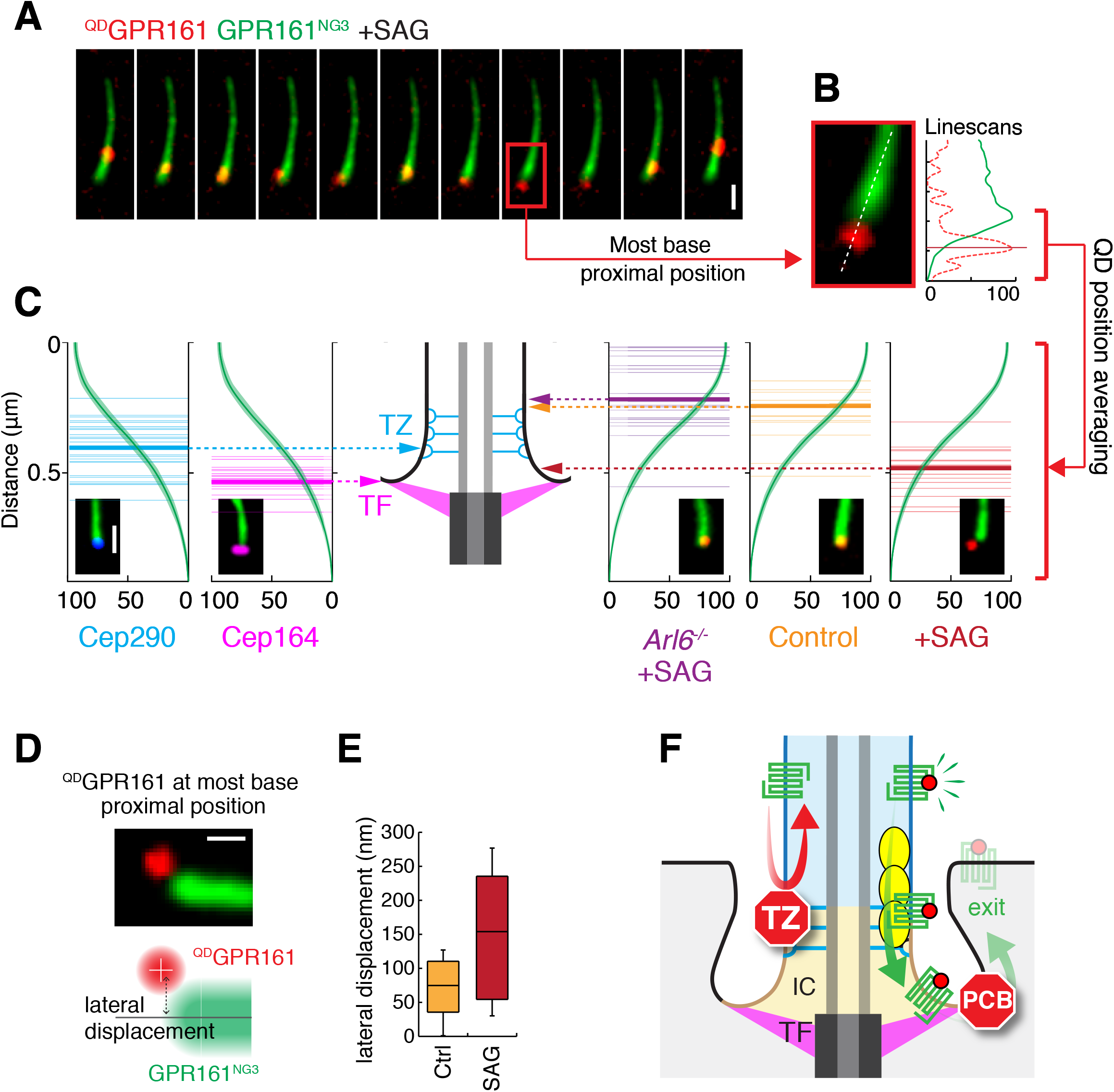
GPR161 traverses the transition zone in an Arl6- and signaling-dependent manner (**A**) Representative images of a base residence event for ^QD^GPR161 after 40 min of SAG treatment. The most base-proximal position of ^QD^GPR161 (red) relative to bulk ^AP^GPR161^NG3^ fluorescence (green) is highlighted by a red box and the enlarged image is shown in (**B**). Scale bar: 1 μm. (**B**) Linescans of the fluorescence intensities of ^QD^GPR161 (red dotted line) and ^AP^GPR161^NG3^ (green line) along the length of cilia. (**C**) Since the longitudinal profile of ^AP^GPR161^NG3^ fluorescence is highly reproducible and unchanged by fixation (Fig. **S5C**), the profile of ^AP^GPR161^NG3^ can be used as a common reference to align the positions of ^QD^GPR161, Cep290 and Cep164 with respect to one another. WT or ***Arl6^-/-^*** IMCD3-[pCrys-^AP^GPR161^3NG^] cells treated with SAG or vehicle for 40 min before imaging. The most base-proximal positions of the centroid of ^QD^GPR161 relative to the profile of ^AP^GPR161^NG3^ were plotted as thin lines and the means plotted as thick lines. The same method was used to plot the positions of immunofluorescence-stained Cep290 and Cep164 relative to ^AP^GPR161^NG3^ fluorescence. Scale bar: 1 μm. ***N*** = 15-26. (**D-E**) The lateral displacement between the centroid of ^QD^GPR161 at its most base-proximal position and the center axis of ^AP^GPR161^NG3^ fluorescence was box plotted. The lateral displacement informs the half-width of the cilium base (control) or the intermediate compartment (SAG). IMCD3-[pCrys-^AP^GPR161^3NG^] cells were treated with SAG or vehicle for 40 min before imaging. Scale bar: 0.5 μm. ***N*** = 11-16 cilia. (**F**) Two-barrier model for exit from cilia. The intermediate compartment is displayed in tan color, the cilium shaft is blue and the cell is gray. GPCR is green, agonist is red, BBSome/Arl6 coats are yellow. TF: transition fibers, TZ: transition zone, IC: intermediate compartment, PCB: periciliary barrier. The diagram is not drawn to scale.

Tracking ^QD^GPR161 over a combined 70 min period of imaging in the presence of SAG revealed that the probability of ^QD^GPR161 entering the intermediate compartment is 0.66 during a one minute interval (Fig. **S5D**). Meanwhile, the absolute exit rate (Fig. **1G**) of 0.0043 molecules/minute is equivalent to the probability of a single molecule of GPR161 experiencing exit during a one minute interval. Thus, a comparison of intermediate compartment entry frequency and ciliary exit rates revealed that less than 1% of intermediate compartment visits productively lead to exit. Meanwhile 99.3% of intermediate compartment visits are resolved by the GPCR returning into the ciliary compartment (Fig. **9A-B, i-ii**). Thus, while crossing the first diffusion barrier at the transition zone appears to be a prerequisite for ciliary exit, it is not sufficient to commit GPCRs for exit because a periciliary barrier blocks lateral diffusion between intermediate compartment and plasma membrane (Fig. **10F**).

Examining the lateral displacement of ^QD^GPR161 to the center of the axoneme when ^QD^GPR161 reached its most base-proximal position in unstimulated cells confirmed that GPR161 stayed within the expected 250 nm diameter of the ciliary membrane during rapid turnaround events in unstimulated cells (Fig. **10D-E**). The mean and maximal lateral displacements of ^QD^GPR161 doubled when ^QD^GPR161 reached its most base-proximal position in SAG-treated cells (Fig. **10D-E**). Since ^QD^GPR161 visits its most base-proximal location in SAG-treated cells while residing within the intermediate compartment, these data indicate that the diameter of the intermediate compartment is close to 550 nm, similar to the 450 nm diameter defined by the tip of the transition fibers (Lau et al., 2012; Yang et al., 2015; Kanie et al., 2017; Yang et al., 2017). The periciliary barrier thus appears to be located near the point where the transition fibers attach to the ciliary membrane.

### Discussion

Our pharmacological and live cell imaging manipulations indicate that, downstream of SSTR3 and Smoothened, a Gαi-mediated decrease in PKA activity promotes BBSome tip redistribution and subsequent retrieval of SSTR3 and GPR161. Intriguingly, the Gα_s_-coupled GPCR Dopamine receptor 1 (DRD1) becomes enriched in cilia of amygdala neurons when BBSome function is compromised (Domire et al., 2011). It will be important for future studies to determine how Gα_s_-coupled GPCRs are retrieved from cilia and whether BBSome tipping is induced by DRD1 activation.

We note that in olfactory receptor neurons, BBS4 is present in all IFT88-marked trains (Williams et al., 2014). In contrast, IFT tracks in unstimulated IMCD3 cells are often devoid of BBSome and we observed frequent uncoupling of BBSome from IFT trains (Fig. **6B**). This suggests that olfactory receptor neurons resemble IMCD3 cells under signaling conditions with respect to BBSome transport. It this context, it is notable that BBS2 and BBS4 accumulate at the tip of olfactory cilia (Williams et al., 2014) and we propose that BBSome transport is highly active in olfactory receptor neurons. According to our absolute quantitation, an average retrograde train contains 10 BBSomes and 62 IFT-B complexes in untreated cells (Fig. **6D** and **G**). The BBSome?IFT stoichiometry (6.2:1) we measure by quantitative imaging is in close concordance with that measured using quantitative mass spectrometry in *Chlamydomonas* (6.5:1) (Lechtreck et al., 2009), thus suggesting a low basal rate of BBSome-mediated transport in *Chlamydomonas* under vegetative conditions.

We find that base confinement events precede irreversible exit from the ciliary compartment. While activation of GPR161 and SSTR3 increased their base confinement frequencies, previous work showed that activation of Smoothened decreases its base confinement frequency (Milenkovic et al., 2015). Together with the accumulation of Smoothened in cilia of unstimulated *bbs* mutant cells (Zhang et al., 2015; Eguether et al., 2014; Goetz et al., 2017), this suggests that Smoothened is constitutively retrieved from cilia by the BBSome and that a reduction in Smoothened retrieval underlies the signal-dependent accumulation of Smoothened in cilia.

Although the idea of a periciliary diffusion barrier was initially considered (Nachury et al., 2010), the transition zone has come to be viewed as the sole diffusion barrier of the cilium in recent years (Garcia-Gonzalo and Reiter, 2012; Gonçalves and Pelletier, 2017; Jensen and Leroux, 2017). Our finding that the ciliary membrane is individualized from the plasma membrane by two successive diffusion barriers suggests the existence of an intermediate compartment located between the transition zone and a nearly impassable periciliary barrier. Crossing of the second barrier is extremely infrequent and is often preceded by a near-complete immobilization prior to exit. In all the exit events we imaged, ^QD^GPR161 diffuses within the plane of imaging after exit, suggesting that the ^QD^GPR161 stayed in the plasma membrane. However, we cannot rule out that this mobility corresponds to an endosome and that endocytosis mediates crossing of the second barrier.

The periciliary barrier is likely to correspond to the recently described Distal Appendage Matrix (DAM) because depletion of the DAM component FBF1 results in the leakage of GPCRs from cilia (Yang et al., 2017). Remarkably, when the receptors for insulin-like growth factor 1 (IGF1) or for transforming growth factor β (TGF-β) undergo signal-dependent exit from cilia, they transiently localize to a zone at the base of cilia that does not overlap with from axonemal markers and may correspond to the intermediate compartment. Furthermore, residence of activated IGF1R and TGF-βR at the base of cilia appears to organize downstream signaling for these two pathways (Clement et al., 2013; Yeh et al., 2013). The intermediate compartment may harbor specific lipids as both PI(4,5)P_2_ and PI(3,4,5)P_3_ are dynamically enriched at a zone at the base of cilia that is clearly non-overlapping from axonemal markers in mammalian cells (Dyson et al., 2017). The intermediate compartment may therefore constitute a privileged signaling locale. Finally, the existence of a second barrier explains why transition zone mutants have only mild mislocalization phenotypes and can still assemble cilia.

The ultrastructural location of the diffusion barrier within the transition zone is beyond the resolution of our imaging study. However, considering that ^QD^GPR161 explores a compartment that is ~ 220 nm long during base confinement events (Fig. **9C**), and given that the distance between the tip of the distal appendages (marked by Cep164) and the proximal end of the transition zone (marked by Cep290) is ~ 100 nm (Yang et al., 2015), it is likely that the intermediate compartment encompasses part of the transition zone and that the diffusion barrier is located within the most distal part of the transition zone (Fig. **10F**).

Furthermore, since activated GPR161 only crosses the transition zone when BBSome/Arl6 coat assembly is permitted, the hypothesis that Hedgehog signaling loosens the diffusion barrier of the transition zone (Dyson et al., 2017) cannot account for GPR161 exit. Instead, our data suggests that BBSome/Arl6 coats bound to retrograde IFT trains on the axoneme-facing side and to cargoes on the membrane-facing side facilitate lateral transport through the transition zone. Thus, in contrast to all other known diffusion barriers, the transition zone is a porous barrier that allows the selective permeation of GPCRs bound to BBSome/Arl6 coats. Physical and genetic interactions between the BBSome and transition zone proteins such as Cep290 (Yee et al., 2015; Goetz et al., 2017; Zhang et al., 2014; Barbelanne et al., 2015) are in agreement with a general model where BBSome/Arl6 coats contact the transition zone during crossing. Taking our data into account, we propose that Arl6-GTP increases the coupling between cargoes, BBSome and IFT trains to facilitate lateral transport through the transition zone. However, the intimate details of transition zone crossing remain to be determined. In particular, the lack of precedent for selective permeation though a membrane diffusion barrier points to distinguishing biophysical features of the transition zone whose definition promises to enrich the concepts underlying diffusion barriers.

## Materials and Methods

### Cell line construction

For all experiments, a mouse inner medullar collecting duct IMCD3-FlpIn cell line was used (gift from Peter K. Jackson). IMCD3-FlpIn cells were cultured in DMEM/F12 (Cat. #11330-057, Gibco) supplemented with 5% FBS, 100 U/mL penicillin-streptomycin, and 2 mM L-glutamine.

Cell lines expressing SSTR3, GPR161, BBS1, BBS5, and IFT88 were generated using the FlpIn System (ThermoFisher Scientific). Construction of multiple expression cassettes with low-expression promoters was conducted as described (Nager et al., 2017). Coding sequences were amplified from plasmids encoding human BBS1 and BBS5 (gifts from Val Sheffield), BirA-ER (gift from Alice Ting (Howarth and Ting, 2008), Addgene plasmid # 20856), mouse GPR161 (BC028163, MGC, Dharmacon), human NPY2R (BC075052, MGC, Dharmacon), mouse IFT88 (I0M20300, UltimateORF, Invitrogen), human MCHR1 (BC001736, MGC, Dharmacon), PACT (Pericentrin and AKAP450 centrosome-targeting domain, gift from Sean Munro (Gillingham and Munro, 2000)), mouse Kif7 (gift from Kathryn Anderson (He et al., 2014, 7)), and mouse SSTR3 (gift from Kirk Mykytyn). BBS1, BBS5, and IFT88 were expressed by the EF1α promoter; SSTR3, NPY2R and MCHR1 the EF1α^Δ^ promoter, and GPR161 by the Crys promoter. Green Fluorescent Protein (GFP), NeonGreen (NG) (Shaner et al., 2013), mScarlet (Bindels et al., 2017) (Addgene #85042), and TandemTomato (tdTomato) (Gift from Michael Davidson (Shaner et al., 2004), Addgene #54653) and an acceptor peptide (GLNDIFEAQKIEWHE) for the biotin ligase BirA (AP) were used in fusion proteins. Kif7 cDNA was subcloned into a modified pCMV-based plasmid (pmScarlet-C, Addgene #85042) that was transfected into IMCD3 cells using Lipofectamine 2000, and clones were selected using Neomycin resistance. The expression level of ^NG3^BBS5 and ^NG3^IFT88 relative to endogenous BBS5 and IFT88 were determined by western blotting (Fig. **3A** and **S2I**), the expression levels of exogenous SSTR3 or GPR161 compared to endogenous GPCRs were determined by measuring the intensity of ciliary fluorescence after immunostaining (Fig. **1C** and **S1B**).

CRISPR-based genome editing was done by transiently expressing Cas9 and guide RNAs (pX330, Addgene #42230, Feng Zhang). Knockouts of *Arl6* and *Ift27* are described (Liew et al., 2014), the guide sequences used were AAGCCGCGATATGGGCTTGC for *Arl6* and GGAAATGGGTCCCGTCGCTG for *Ift27.* Knockouts of *Arrb1* and *Arrb2* are described (Nager et al., 2017), the guide sequences used were ACTCACCCACGGGGTCCACG for *Arrb1* and TCTAGGCAAACTTACCCACA for *Arrb2..* To generate a *Tulp3^-/-^* cell line, a guide RNA targeting the sequence ACGTCGCTGCGAGGCATCTG was used. CRISPR-modified clones were isolated by limited dilution, and verified by western blotting for the disrupted gene product. To confirm gene editing, the modified genomic locus was isolated used using the ThermoFisher CloneJET PCR cloning kit (Cat. #K1231, ThermoFisher Scientific).

### Low Expression Promoters

Cloning of constructs with low expression promoters was done as described (Nager et al., 2017). Briefly, an *NsiI* restriction cloning site was inserted by site directed mutagenesis before the EF1α promoter in pEF5B/FRT (Nager et al., 2017). The EF1α promoter was then excised by NsiI and SpeI, and replaced by either the UbC, thymidine kinase (TK), EF1α^Δ^, CMV^Δ^, or a minimal chicken lens δ-crystallin promoter (Morita et al., 2012; Ferreira et al., 2011; Kamachi and Kondoh, 1993). The UbC promoter was cloned from pLenti6/UbC/V5-Dest (Cat. #V49910, ThermoFisher Scientific). The thymidine kinase promoter was from pRL TK (Cat. #E2241, Promega). EF1α^Δ^ consists of a TATA-less EF1α promoter from pEF5/FRT/V5-Dest (Cat. #V602020, ThermoFisher Scientific) wherein the TATA box sequence, TATAA, was mutated to TCCCC. CMV^Δ^ was designed after the CMV(Δ6) promoter (Morita et al., 2012), and was cloned from pcDNA3.1 (Cat. #V79020, ThermoFisher Scientific). The chicken lens δ-crystallin promoter was cloned from a pGL3-8xGli-Firefly-Luciferase plasmid (gift from Phil Beachy).

### Hippocampal Neurons

Rat hippocampal neurons were dissected from postnatal day 0 or 1 rat pups and plated on poly-D-lysine-coated 12 mm #0 cover glass. Neurons were cultured in Neurobasal medium with serum-free B27 (Cat. #21103049, ThermoFisher Scientific) and Gibco GlutaMAX (Cat. #35050061, ThermoFisher Scientific). Neurons were identified by nuclear NeuN staining in immunofluorescence studies.

### Transfection

Plasmids were reverse-transfected by Lipofectamine 2000 (Cat. #11668027, ThermoFisher Scientific). Briefly, detached cells were plated with the transfection reagent and plasmid in Optimem (Cat. #31985070, ThermoFisher Scientific). The transfection reagent was replaced by fresh DMEM/F12 medium after 4 hr. siRNAs were reverse-transfected by Lipofectamine RNAiMAX (Cat. # 13778030, ThermoFisher Scientific). Briefly, detached cells were plated with the transfection reagent and the siRNA duplex in Optimem. Cells were then grown for 48 h before 24 h starvation and subsequent imaging. Control (Cat. #D-001210-04-05), BBS1 (Cat. #D-019180-03), BBS2 (Cat. #D-010080-02) and BBS4 (Cat. #D-054691-03) were from Dharmacon, and the Kif7 siRNA (Cat. #GS16576) and matched control (Cat. #1027280) were from Qiagen. siRNA targeting Arl6 (CTTTAGGACTTGAGACATT) was described (Jin et al., 2010).

### Pharmacology

Small molecules were added at the following concentrations and for the indicated pretreatment times unless otherwise indicated: ACQ090 (20 μM, gift from Novartis (Bänziger et al., 2003)), IBMX (500 μM, 30 min pre-incubation, Cat. # I5879, Sigma-Aldrich), L796,778 (10 μM, gift from Merck (Rohrer et al., 1998),), PKI (50 μM, 40 min pre-incubation, Cat. BML-P210-0500, Enzo Life Sciences), PTX (10 ng/mL, 16 h pre-incubation, Cat. #180, List Biological Laboratories), Rp-cAMPS (10 μM, 2 h pre-incubation, Cat. #sc-24010, Santa Cruz), SAG (200 nM, ALX-270-426-M001, Enzo Life Sciences), Somatostatin (10 μM, Cat.#ASR-003, Alomone Labs), SQ22536 (500 μM, 40 min pre-incubation. Cat. #S153, Sigma-Aldrich).

### Antibodies and Affinity Reagents

The following antibodies were used: AC3 (sc-588, Santa Cruz Biotechnology), acetylated tubulin (Clone 6-11B-1, Sigma-Aldrich), Arl6 (Jin et al., 2010), ß-Arrestin1 (Cat. #15361-1-AP, Proteintech Group), β-Arrestin2 (Cat. #10171-1-AP, Proteintech Group), BBS5 (Cat. #14569-1-AP, Proteintech Group), BBS9 (Cat. #HPA021289, Sigma-Aldrich), Centrin (Clone 20H5, Millipore), Cep164 (gift from Tim Stearns), Cep290 (gift from Sophie Saunier), c-Myc (Cat. #sc-40, Santa Cruz Biotechnology), GFP (Cat. #A11122, Invitrogen), GPR161 (gift from Saikat Mukhopadhyay), IFT139 (gift from Pamela Tran), IFT140 (Cat. #17460-1-AP, Proteintech Group), NeuN (Cat. #MAB377, Millipore), Ninein (gift from Michel Bornens), Phospho-p44/42 MAPK Erk1/2 Thr202/Tyr204 (Cat. #4370, Cell Signaling Technologies), SSTR3 (Cat. #sc-11617, Santa Cruz Biotechnology), and TULP3 (gift from John Eggenschwiler). Biotinylated SSTR3 and GPR161 were detected using Alexa647-labeled monovalent streptavidin (Ye et al., 2013).

### Protein-Protein Interaction Assays

IFT-A was purified through a LAP tag on the N-terminus of IFT43 as described (Nachury, 2008). Briefly, IMCD3-[pEF1α-^LAP^IFT43] were harvested and lysed, and the IFT-A complex was captured by GFP immunoaffinity and eluted with the TEV protease. GST and GST-SSTR^i3^ fusion proteins were purified from *E.coli,* and used for interaction assays as described (Jin et al., 2010).

The interaction of ^GFP^Kif7 with ^Myc^BBS1 was assayed by co-transfection/co-IP. HEK293 cells were forward transfected in a 6-cm plate with plasmids expressing ^GFP^Kif7 and either MycRFp or a Myc-tagged BBSome subunit. After two days, cells were washed and treated with media containing either IBMX or DMSO for 30 min. Cells were then trypsinized, pelleted and cleaned in a flacon tube, and lysed for 10 min with cold CoIP buffer (50 mM Tris pH 7.4, 150 mM NaCl, 1% Triton X-100, 1 mM DTT, 1 mM AEBSP, 800 nM Aprotonin, 15 μM E-64, 10 mg/ml Leupeptin, 10 mg/ml Pepstatin A, 10 mg/ml Bestatin). The resulting lysates were then centrifuged for 15 min, concentration-matched, and then added to anti-GFP antibody coupled beads in CoIP buffer. The beads were rotated for 20 min at 4^o^C, pelleted and washed with CoIP buffer 4 times, and eluted by boiling in SDS-PAGE loading buffer.

### Absolute quantitation of cilia-localized proteins

In Fig. **1C**, the number of ^AP^SSTR3^GFP^ molecules per cilium resulting from expression driven by the EF1α, UBC, TK, EF1α^Δ^, and CMV^Δ^ promoters was estimated by comparison to a viral particle containing exactly 120 GFP molecules (Breslow et al., 2013). As ciliary GFP was not detectable by direct imaging of the IMCD3-[pCrys-^AP^SSTR3^GFP^] line, immunostaining for GFP was used to amplify the fluorescent signal and compare to immunostained GFP in IMCD3-[pEF1α^Δ^-SSTR3^GFP^] cells. To estimate the number of SSTR3 molecules per hippocampal neuron, hippocampal neurons and IMCD3-[^AP^SSTR3^GFP^] cells using the aforementioned promoters were compared by SSTR3 immunostaining. The epitope recognized by the SSTR3 antibody (Cat. #sc-11617, Santa Cruz Biotechnology) is identical between rat and mouse (Santa Cruz Biotechnology, personal communication).

For quantitation of NeonGreen-tagged proteins including ^AP^SSTR3^NG^, ^AP^GPR161^NG3^, ^NG3^BBS5, and ^NG3^IFT88, ciliary molecules were quantified using single NeonGreen trimers (NG3) as a calibrator (Prevo et al., 2015). NG3 protein was recombinantly expressed in *E. coli,* and purified by nickel affinity. The purified protein was then sparsely immobilized on a 18x18 mm coverslip for imaging on a DeltaVision microscope. Imaging was done in Invitrogen Live Cell Imaging Solution (Cat. #A14291DJ). To confirm that fluorescent foci originate from a single NG3, we confirmed that fluorescence was lost by photobleaching in three discrete steps (Fig. **1E**, **S1H**). The mean fluorescent intensity from over 1257 molecules was then used to estimate the number of NeonGreen-tagged molecules in cilia.

To quantify the absolute number of ciliary proteins, serum staved live IMCD3 cells were imaged in Invitrogen Live Cell Imaging Solution (Cat. #A14291DJ) with the same exposure setting as used in single NG3 fluorescent quantitation. Therefore,

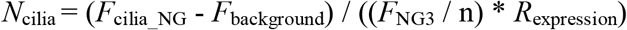

Where N_cilia_ is the absolute number of ciliary protein, F_cilia___NG_ is the total ciliary fluorescence detected from NG or NG3 labeled proteins, F_background_ is the background fluorescence measured in the adjacent area. F_NG3_ is the fluorescent intensity of a single NG3 protein, n determined by the single or triple NG tag was used. For example, n = 3 for ^AP^SSTR3^NG^ and n=1 for GPR161^NG3^. R_expression_ is the abundance ratio between the NeonGreen-tagged form of a given protein and the total amount of that protein in cilia. For pEF1α-^NG3^BBS5 and pEF1α-^NG3^IFT88, R_expression_ = 0.55 and 0.51 respectively, which were measured by western blotting (Fig. **3A** and **S2I**). For pEF1α^Δ^-^AP^SSTR3^NG^, *R*_expression_ = 1 since IMCD3 cells do not express SSTR3. For ^AP^GPR161^NG3^, *R*_expression_ could not be directly measured because tagging of GPR161 at the C-terminus (e.g. in GPR161^NG3^) interferes with recognition by the anti-GPR161 antibody developed by Mukhopadhyay and colleagues (Mukhopadhyay et al., 2013). We thus utilized ^AP^GPR161 as an intermediate calibrator between ^AP^GPR161^NG3^ and endogenous GPR161. A plasmid encoding pCrys-^AP^GPR161 was transfected into IMCD3 cells and the relative expression levels of transiently expressed ^AP^GPR161 and stably expressed ^AP^GPR161^NG3^ were measured by mSA647 pulse-labeling.

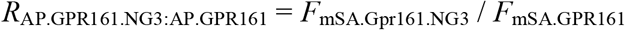

Where R_AP.GPR161.NG3:AP.GPR161_ is the ratio between stably expressed ^AP^GPR161^NG3^ and transiently expressed ^AP^GPR161 in cilia, F_mSA GPR161_ is the ciliary fluorescence signal measured by mSA647 labeling of transiently expressed ^AP^GPR161, and F_mSA.GPR161.NG_3 is the ciliary fluorescence signal measured by mSA647 labeling of stably expressed ^AP^GPR161.

The relative ciliary expression levels of endogenous GPR161 and ^AP^GPR161 were measured by anti-GPR161 antibody.

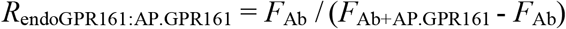

Where R_endoGPR161: AP.GPR161_ is the ratio between endogenous GPR161 and transiently expressed ^AP^GPR161 in cilia, *F*_Ab_ is the ciliary fluorescence signal measured by immunofluorescence of untransfected cells with the anti-GPR161 antibody and *F*_Ab+AP.GPR161_ is the ciliary fluorescence signal measured by immunofluorescence of cells transiently expressing with ^AP^GPR161 with the anti-GPR161 antibody.

Therefore,

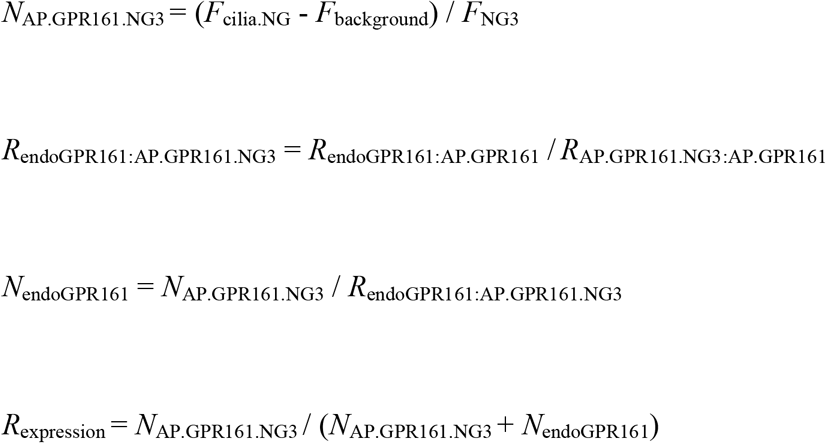

Where *R*_endoGPR161·AP.GPR161.NG3_ is the ciliary abundance ratio between endogenous GPR161 and stably expressed ^AP^GPR161^NG3^; *N*_endoGPR161_ and *N*_AP.GPR161.NG3_ are the absolute number of endogenous GPR161 and stably expressed ^AP^GPR161^NG3^ in cilia; *F*_cilia___NG_ is the total ciliary fluorescence detected in the green channel from ^AP^GPR161^NG3^; *F*_background_ is the background fluorescence measured in the adjacent area; *F*_NG3_ is the fluorescent intensity of a single NG3 protein; *R*_expression_ is the ciliary abundance ratio between stably expressed ^AP^GPR161^NG3^ and total GPR161. Using the above strategy, we determined that:

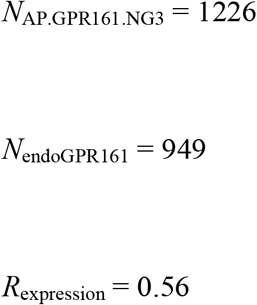

### Bulk GPCR Exit Assays

Bulk measurements of GPCR exit were done as described previously (Nager et al., 2017). Briefly, to measure SSTR3 exit, cell expressing ^AP^SSTR3^NG^ were firstly washed three times with PBS containing 5 mM MgCl_2_ (PBS-Mg) and then pulse-labeled with Alexa647-labeled mSA (mSA647) for 5-10 min. To remove the unbound mSA647, cells were washed three times with PBS containing 5 mM MgCl_2_ and imaged on a DeltaVision microscope following addition of somatostatin. For each time point, the integrated Alexa647 fluorescence density was measured using ImageJ. The cilia-adjacent fluorescence was subtracted as the background, and a mathematical photobleaching correction was applied:

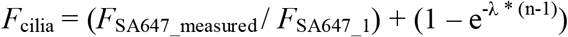

Where λ is the photobleaching decay constant, n is the number of images taken, F_SA647_measured_ is the integrated SA647 fluorescence measured for image ‘n’, *F*_SA657_1_ is the measurement for the first time point, and *F*_cilia_ is the reported fluorescence. In this equation, *F*_cilia_ is reported in Relative Fluorescence Units (RFUs).

^AP^GPR161^3NG^ was assayed similarly except that quantitations were done by NG fluorescence intensity using the following equation:

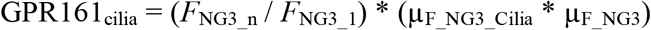

Where GPR161_cilia_ is the number of GPR161 molecules in the cilium, *F*_NG3_n_ is the photobleaching-corrected integrated NG fluorescence measured for image ‘n’, F_NG3_1_ is the measurement for the first time point, μ_F_NG3_cilia_ is the mean integrated NG fluorescence from a population of ^AP^GPR161^NG3^ cilia, and μ_F_NG3_ is the mean integrated green fluorescence from individual NG3 molecules.

To measure the exit rate of SSTR3 and GPR161 in the first 2h after adding agonist (Fig **1F-G**),

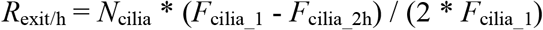

Where *R*_exit/h_ is the number of SSTR3 or GPR161 that exited cilia per hour, *F*_cilia_1_ is the background corrected ciliary fluorescent intensity before adding agonist, *F*_cilia_2_h is the background corrected ciliary fluorescent intensity after 2h of agonist treatment, *N*_cilia_ is the absolute number of SSTR3 or GPR161 molecules in cilia as described in previous section.

To assess significant differences in GPCR removal, we used multiple regression. First, raw data was linearly fitted (F_cilia_ = m * time + c). Conditions were then compared using a z-statistic:

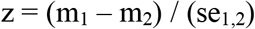

Where m_1_ and m_2_ are the fitted slopes for two experiments, and se_1,2_ is the propagated standard error of the slopes:

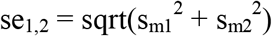

Where s_m1_ and s_m2_ are the standard deviations for m_1_ and m_2_, respectively. Z-statistics were converted to p-values for statistical interpretation.

### Fixed Imaging

In a 24-well plate, 50,000 to 100,000 cells were seeded on Fisherbrand acid-washed cover glass (12 mm #1.5, Cat. #12-545-81, Fisher Scientific). Cells were grown for 24 h, and then starved for 16-24 h in 0.2% FBS media prior to experimental treatment. After treatment, cells were fixed with room-temperature 4% paraformaldehyde in phosphate buffer saline (PBS) for 5 min, extracted in −20°C methanol for 5 min, permeabilized in PBS containing 0.1% Triton-X100, 5% normal donkey serum (017-000-121, Jackson Immunoresearch Labs), and 3% bovine serum albumin (BP1605-100, ThermoFisher Scientific) for 30 min, and subsequently immunolabeled for imaging. Briefly, cells were incubated with primary antibodies for 1 h, washed three times with PBS, incubated with dye-coupled secondary antibodies (Jackson Immunoresearch Labs) for 30 min, washed two times with PBS, stained with Hoechst DNA dye, washed twice more with PBS, and mounted on slides using Fluoromount-G (Cat. #17984-25, Electron Microscopy Sciences). Cells were then imaged on a DeltaVision system. In most experiments, cilia closest to the cover slip were imaged (ventral cilia) as these cilia often lay perpendicular to the objective and within a single focal plan. In select cases, cilia pointing away from the coverslip (dorsal cilia) were imaged as to reduce background fluorescence. To do so, a Z-stack of images with 0.3-μm separation was collected and deconvolved using SoftWoRx 6.0.

### Live-Cell Imaging

400,000 cells were seeded on acid-washed 25 mm cover glass (Cat. #72223, Electron Microscopy Sciences) in a 6 cm dish. After 24 h of growth, cells were starved for 16 h and transferred to the DeltaVision stage for imaging at 37 °C inside an environmental chamber. Cells were imaged in DMEM/F12 media, with HEPES and no phenol red and 0.2% FBS (Cat. #11039-021, Gibco). For all >1 h imaging experiments, the imaging chamber was overlaid with a petri dish containing a moist towel to maintain the imaging volume and pH. To accurately measure removal of ^AP^SSTR3^NG^, the biotinylated AP tag of SSTR3 was pulse-labeled with Alexa647-labeled monovalent streptavidin (SA647) for 5 min as described (Ye et al., 2013).

All imaging was conducted on an Applied Precision DeltaVision equipped with a PlanApo 60×/1.40 numerical aperture (NA) oil objective lens and a PlanApo 60×/1.49 NA total internal reflection microscopy (TIRF) oil objective lens (Olympus) and a 488 nm laser from DeltaVision Quantifiable Laser Module (QLM), and images were captured with a pco.edge sCMOS camera (PCO) with near-perfect linearity across its 15 bit dynamic range. The pixel size of the sCMOS camera is 0.1077 μm.

### Kymograph Analysis and Processivity

To analyze intraciliary trafficking, IFT88 and BBS5 proteins were genetically labeled with NG3 and rapidly imaged with TIRF (4 Hz) for short time periods (30-60 s). TIRF illumination reduced nonciliary (background) fluorescence, permitting visualization of dim trains. The resulting movies were analyzed by ImageJ for generating kymographs. KymographClear and KymographDirect (Mangeol et al., 2016) were used to deconvolve anterograde from retrograde trains for measuring relative intensities for BBSome trains whereas Multi Kymograph was used for unaltered presentation (e.g. Fig. **6A**).

The processivity of IFT-B or BBSome trains along axonemal microtubules was defined as the duration for which a fluorescent focus of IFT88 or BBS5 unidirectionally moved along the the cilium at a rate expected for kinesin or dynein-mediated transport (0.3-0.6 μm/s). Only events occurring for longer than 3 seconds were considered. Measurements were made from kymographs of ^NG3^IFT88, ^NG3^BBS5, or ^tdTomato^IFT88 and ^NG3^BBS5. Briefly, kymographs were visualized in ImageJ, and lines were drawn along the long axis of the cilium (“the leg”) and the processive movement (“the hypotenuse”). The angle between the leg and the hypotenuse was measured (“the included angle”). Comparison of the leg and the hypotenuse was used to quantify the processivity of a IFT or BBS-containing train. Comparison of the hypotenuse and the included angle was used to measure the velocity of IFT and BBS trains.

### Quantitation of IFT/BBS trains

To estimate the number of IFT88 or BBS5 molecules per train, the total NG3 fluorescence along the ciliary axoneme was first divided by the number of trains:

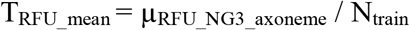

Where T_RFU_mean_ is the estimated ^NG3^IFT88 or ^NG3^BBS5 fluorescence of a single train, μ _RFU_NG3_axoneme_ is the mean integrated fluorescence along the axonemes of several cilia (excluding the base and the tip), and N_train_ is the number of trains counted by kymograph analysis in the cilia when μ_RFU_NG3_axoneme_ was measured. To increase measurement precision for calculating μ_RFU_NG3_axoneme_, cilia were imaged by epifluorescence rather than TIRF illumination as the TIRF field did not reproducibly illuminate cilia from different cells. T_RFU_mean_ was then used to calculate the absolute number of IFT88 or BBS5 molecules per train by using the NG3 standard described above (see “Absolute quantitation of cilia-localized markers”):

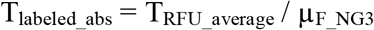

Where T_labeled_abs_ is the average number of labeled molecules per train, and μ_F_NG3_ is the previously-measured mean fluorescence of a single NG3. As T_labeled_abs_ does not account for unlabeled IFT88 or BBS5 within a train, a correction must be applied that relates the ratio of unlabeled to labeled molecules in the cell:

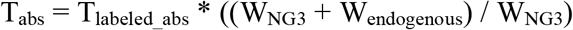

Where T_abs_ is the number of IFT88 or BBS5 molecules per train, and W_NG3_ and W_end_o_g_eno_us_ are the integrated intensities of western blot bands for NG3-tagged and endogenous molecules using antibodies against mouse IFT88 or BBS5 (Fig. **3A** and **S2I**). T_abs_ was used to calculate the number of molecules in an anterograde (T_abs_anterograde_) versus retrograde (T_abs_anterograde_) train:

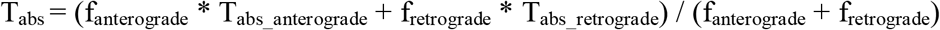

Where f_anterograde_ and f_retrograde_ are the frequency of either anterograde or retrograde trains. Using KymographClear analysis on ^NG3^IFT88, we measure f_anterograde_ = 21.4 trains/min and fretrograde = 18.5 trains/min. Tabs_anterograde and Tabs_retrograde can now be solved using the size ratio (RT) of anterograde versus retrograde trains:

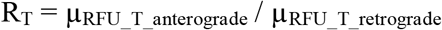

Where μ_RFU_T_anterograde_ and μ_RFU_T_retrograde_ are the mean NG3 intensity of anterograde or retrograde trains. By KymographClear analyses, R_T_ = 1.69 for ^NG3^IFT88, and R_T_ = 1.78 for ^NG3^BBS5. Relating T_abs_anterograde_ to T_abs_retrograde_:

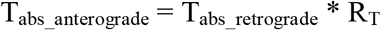

One can calculate the number of IFT88/BBS5 particles in anterograde and retrograde trains:

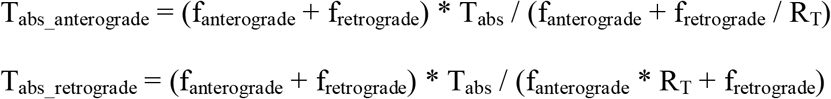

### Linescan and tipping quantitation

The longitudinal fluorescence intensities of BBS5, IFT88, SSTR3 and GPR161 were measured in ImageJ by a Plot Profile of a 5-pixel-wide line along the long axis of the cilium. To average data from multiple cilia, pixel Intensities were assigned a length-percent with 0% referring to the base, and 100% referring to the tip. Values were then grouped into 5% bins and averaged. Bin means were then averaged across multiple cilia and plotted (e.g. Fig. **3B**).

To quantify ^NG3^BBS5 tipping, a 5x7 pixel box was centered at the cilium tip, and the integrated fluorescence intensity within the box was measured by ImageJ. The resulting values were background corrected by subtracting the fluorescence immediately adjacent to the cilium. The values from multiple ciliary tips were averaged, and then converted to absolute numbers of BBS5 particles using the NG3 calibration.

### Single-molecule labeling with ^SA^Qdot

Molecules of SSTR3 or GPR161 present on the surface of IMCD3-[pEF1α^Δ-AP^SSTR3^NG^] and IMCD3-[pCrys-^AP^GPR161^NG3^] cells were labeled with ^SA^Qdot through the affinity of streptavidin for the extracellular biotinylated AP tag. Both cells lines express an ER-targeted BirA ligase that biotinylates the N-terminal AP tag of each GPCR (Nager et al., 2017; Howarth and Ting, 2008). Single GPCR labeling was achieved through a blocking strategy. First, to block surface-exposed biotinylated receptors, cells were washed with PBS (3 times), and then incubated with 10 nM mSA for 15 min. Cells were washed with PBS (3 times), and incubated with 1 nM ^SA^Qdot (Cat. #Q10123MP, ThermoFisher Scientific) solution for 15 min. As the ^SA^Qdot only binds to receptors newly arrived at the surface during the 15 min of labeling, this approach labels very few receptors. Lastly, cells were washed with PBS (3 times), and transferred to imaging media (DMEM/F12, HEPES, no phenol red media; Cat. #11039-021, Gibco) with 0.2% FBS and 1 μM biotin to block unbound Streptavidin on ^SA^Qdot.

Qdot-labeled SSTR3 and GPR161 molecules exited the cilium at rates congruent with bulk imaging (Fig. **1F-G**). By counting the number of ^QD^SSTR3 per cilia after 2 h treatment with vehicle or sst (Fig. **8C**), we determined that sst treatment resulted in the loss of 25% of cilia-localized Qdots, consistent with the SSTR3 exit rate measured by bulk fluorescence (Fig. **1F**). As a second test, the frequencies of ^QD^GPR161 exit events captured by live imaging matched the frequencies predicted from imaging the decrease in bulk fluorescence of GPR161^NG3^ (Fig. **S5B**).

### Qdot localization and analysis

To monitor single receptor trafficking, cilia bearing one Qdot were imaged on the Deltavision microscope at 2 Hz for 5 to 20 min. Cilia were identified by the NG signal from the C-terminal NG and NG3 tags. The centroid of ^SA^QD-labeled single SSTR3 and GPR161 were mapped as described (Sage et al., 2005). The localization precision of Qdot was measured as described (Deschout et al., 2014).

Processivity, residence, and confinement frequencies are based on kymograph analyses (examples shown in Fig. **S4**). Processive ^QD^GPCR movements were defined as consecutive frames were a Qdot moved longitudinally along the ciliary axoneme. To distinguish motor-driven from diffusive events, only processive motions that extend for at least 6 consecutive frames (Fig **S3G**) or 10 consecutive frames (Fig. **8F**) are plotted. Residence at the base and tip was operationally defined as residing at the basal or distal 10% length of the cilium for longer than 2.5 seconds (Fig. **8G**). As freely diffusing receptors occasionally remain at the cilia base or tip for more than 2.5 seconds, confinement was defined by a more stringent criterion of events lasting longer than 15 seconds (Fig. **8G**).

The most base-proximal position of ^QD^GPR161 was mapped on a coordinate system established by the profile of ciliary GPR161^NG3^ (Fig. **10A-C**). The same approach was applied to map the immunofluorescence intensity center of Cep290 and Cep164 (Fig. **10A-C**). Intermediate compartment visits were defined as events where the QDot centroid crossed the 50^th^ percentile of NG intensity in the GPR161^NG^ longitudinal scan profile (Fig. **10C and F**).

### Box Plots

All box plots display the second, third, and fourth quartiles along with whiskers that represent values within the 1.5x the interquartile range. Outliers exceeding the whiskers are plotted as points.

### Online supplemental material

Fig. **S1** shows that low-expression promoters recapitulate the physiological exit kinetics of SSTR3. Fig. **S2** identifies IFT-A/Tulp3 as the importer of GPCRs into cilia and BBSome/Arl6 as the exporter. Fig. **S3** presents BBSome and IFT train processivity and single molecule quantitation. Fig. **S4** shows kymographs of single molecules of ^QD^SSTR3 and ^QD^GPR161. Fig. **S5** shows visualization of ^QD^GPR161 exit at single molecule resolution. Movie **S1** shows somatostatin-dependent removal of SSTR3 from cilia. Movie **S2** shows IFT-B foci movements in cilia. Movie **S3** shows co-movement of BBSome and IFT-B. Movie **S4** shows BBSome foci movement in cilia. Movie **S5** shows dynamics of BBSome-mediated retrieval. Movie **S6** shows behavior of Qdot-labeled SSTR3 in control-treated cells. Movie **S7** shows behavior of Qdot-labeled SSTR3 in somatostatin-treated cells. Movie **S8** shows direct observation of ciliary exit of Qdot-labeled GPR161.

## Acknowledgments

We thank M. Bornens for the Ninein antibody, S. Saunier for the Cep290 antibody, T. Stearns for the Cep164 antibody, K. Anderson for the Kif7 cDNA, S. Mukhopadhyay for the GPR161 antibody, P. Beachy for the δ-crystallin promoter, J. Eggenschweiler for the Tulp3 antibody, C. Smith for the BBS5 monoclonal antibody, M. Lin for neurons, T. Gadella for mScarlet cDNA, Novartis for ACQ090, and A. Kopp and J. Goldstein for technical assistance. We thank Drs. K. Hill, A. Benmerah and the Nachury lab for comments on the manuscript. This work was supported by NIH funding (GM089933 to M.V.N) and a Fayez Sarofim/Damon Runyon Fellowship to A.R.N. (DRG 2160-13). The authors declare no competing financial interests.

## Author contributions

M.V.N. conceived and coordinated the project and wrote the paper with contributions from all authors; F.Y. and A.R.N jointly developed the cell lines used for imaging of GPCRs and BBSome; F.Y. conducted all single-molecule imaging and the quantitative analysis of BBSome trains and exit. Data described can be found in the main figures and supplementary materials. The authors declare no conflict of interest.

